# LncRNA Hmrhl regulates expression of cancer related genes in Chronic Myelogenous Leukemia through chromatin association

**DOI:** 10.1101/2020.09.17.301770

**Authors:** Subhendu Roy Choudhury, Sangeeta Dutta, Utsa Bhaduri, Manchanahalli R Satyanarayana Rao

## Abstract

Long non-coding RNA has emerged as a key regulator of myriad gene functions. One such lncRNA mrhl, reported by our group, was found to be a regulator of *SOX8*, Wnt-signalling along with an important role in embryonic development in mouse. Recently, its human homolog, human mrhl (Hmrhl) was uncovered and study revealed its differential expression in several type of cancers, notably leukemia. In the present study, we further characterize molecular features of lncRNA Hmrhl and gain insight into its functional role in leukemia by gene silencing and transcriptome-based studies. Results indicate its high expression in CML patient samples as well as in K562 cell line. Silencing experiments suggest role of Hmrhl in cell proliferation, migration & invasion in K562 cells. RNA-seq and ChiRP-seq data analysis further revealed its association with important biological processes, including perturbed expression of crucial TFs and cancer-related genes. Among them ZIC1, PDGRFβ and TP53 were identified as regulatory targets, with high possibility of triplex formation by Hmrhl at their promoter site. In addition, we also found TAL-1 to be a potential regulator of Hmrhl expression in K562 cells. Thus, we hypothesize that Hmrhl lncRNA may play a significant role in the pathobiology of CML.

## INTRODUCTION

Long non-coding RNA (lncRNA) has emerged as the most studied gene regulatory element of this decade. As the name implies, long non-coding RNAs are transcripts with a length longer than 200 nucleotides, which have no functional protein coding ability whatsoever (1, 2). Reported to be expressed from unicellular eukaryotic organism to humans, lncRNAs show tissue/cell/stage specific differential expression, are not well conserved, exhibit different sub-cellular localization and shows enormous diversity in its role and mechanism to regulate expression of protein coding genes (3–7). With the recent developments in deep sequencing technologies, their repertoire is exponentially increasing, however biological significance and characterization of most of lncRNA remains elusive (8, 9). Some lncRNAs which are well studied are reported to be the key players in distinct processes like dosage compensation, genomic imprinting, epigenetic regulator, pluripotency, post transcription regulator of mRNA, modulator of stability/translation of mRNA (10–16). LncRNAs have thus become the focal point of genomic research as they keep signifying their role as crucial regulator in vital biological processes like development and differentiation, cell cycle progression, and most importantly in pathology and progression of many human diseases including cancer (7, 16–21). There is now enough evidence and knowledge that cellular stage/type specificity of these lncRNAs can provide us better understanding, identification, prognostic value and even therapeutic options for many incurable diseases especially cancer (22–25).

Transcriptome analysis of normal and cancerous tissues over the years has revealed differential expression of at least 2000 lncRNAs, with some been specific to certain type of cancers (26). For example, expression of differential display 3, also known as PCA3 and SCHLAP1 were found to be specific for human prostate cancer with PCA3 been later approved by the Food and Drug Administration (FDA) to be used as a biomarker for prostate cancer (27, 28). SAMMSON which has melanoma-specific expression provides a potential target for therapeutics with fewer side effects (29). Contrary to this, many well-known lncRNAs are reported to be involved in various cancers, such as deregulation of HOTAIR is associated with 26 different human tumors (30). LncRNA, MALAT1 was found to be overexpressed in several human cancers, including lung, breast, prostate, hepatocellular and ovarian cancer while H19 in human cancers, including liver, breast, gastric, colorectal, esophageal and lung cancer (31–33). In addition, differential expression of many lncRNAs like ANRIL, NEAT1, LUNAR1 and PCGEM1 are reported to be associated with several different cancer types (34–37).

LncRNA Mrhl (Meiotic recombination hot spot locus RNA) was first reported in mouse by our group and has been extensively studied since then to reveal its biological significance (38–43). This 2.4kb, intronic, nuclear restricted lncRNA was found to be a negative regulator of Wnt signaling via its interaction with p68/Ddx5 RNA helicase in mouse spermatogonial cells (39, 40). In addition, the role of mrhl in meiotic commitment of spermatogonial cells through regulation of *SOX8* at the chromatin level has been documented (41). More recently, transcriptomics and genome-wide occupancy studies of Mrhl in mouse embryonic stem cells (mESCS) revealed its role as a chromatin regulator of cellular differentiation and development genes along with its probable importance in maintenance of stemness in mESCS (43).

In our latest study, we have identified the human homolog of mouse mrhl (Hmrhl) which shares 65% homology with that of mouse mrhl and is encoded in an identical syntenic locus within the *phkb* gene (44). This study has revealed a functional role of Hmrhl as an enhancer RNA for its host gene, *phkb,* in chronic myelogenous leukemia (CML) cell line. Additionally, expression profile of Hmrhl among various tumor and normal samples confirmed its deregulation in several cancers (44). Among these, significant upregulation of Hmrhl in lymphoma tumor samples was evidently notable. This observation prompted us to gain insight into the possible functional molecular role of Hmrhl in leukemia. We started with the expression and characterization of hmrhl in CML cell line, K562. Our sub-cellular localization study specified its enrichment within the nucleus and its association with chromatin which was consistent with our earlier observation with mouse mrhl RNA. *In-vitro* studies demonstrated that depletion of Hmrhl influences cancer related phenotypes like cell migration, proliferation, and cell invasion in K562 cells. We used RNA-seq based approach to address the effect of silencing Hmrhl on the global transcriptome of K562 cells which revealed deregulation of 831 genes with significant perturbation of developmental processes and enrichment in chemokines and cytokine-mediated signaling pathway. Additionally, genome-wide occupancy study of Hmrl RNA indicated its association with several loci throughout the genome, particularly at the intergenic and repetitive elements region along with other regions. Interestingly, by overlapping data from RNA seq and ChIRP seq, we found possible involvement of Hmrhl in the regulation of *PDGFRβ* and *ZIC1* genes. Observations from our data also suggested that regulation of Hmrhl itself is mediated through TAL1, a key transcription factor that is involved in hematopoiesis. Deductions from this study hence suggest Hmrhl as a chromatin regulator of genes associated with phenotype of leukemia, mechanism of which needs to be explored further.

## MATERIAL AND METHODS

### Cell lines and Reagents

K562 (Chronic Myelogenous Leukemia/ Erythro Leukemia) cell line was obtained from NCCS, Pune (India), whereas human lymphocyte cell line (ID no. GM12878) was from Coriell Institute, USA. Both cell lines were cultured in RPMI medium (Gibco) supplemented with 10% FBS (Gibco) for K562 and 15% non-heat inactivated FBS (Gibco) for GM12878 along with 100 units/ml penicillin-streptomycin solution (Sigma) at 37°C in a humidified chamber with 5% CO_2_. All chemicals were obtained from Sigma-Aldrich and Life Technologies unless otherwise specified.

### RNA isolation, cDNA synthesis and real-time PCR

Total RNA was isolated from K562 and GM12878 cells using RNAiso Plus (Takara Bio) as per the manufacturer’s instructions. RNA samples were further treated with DNase (New England Biolab) to remove any genomic DNA contamination. About 500-1000ng RNA was used as a template for cDNA synthesis using a cDNA synthesis kit from Biorad. The cDNA was further diluted to 1:10 with nuclease-free water, and real-time PCR was carried out using the SyBr green mix (Takara) in a real-time PCR machine (Biorad CFX96 Real-Time System). A pool of four Hmrhl specific siRNAs targeting the conserved region of Hmrhl and scrambled negative control siRNA were obtained from Dharmacon. K562 cells were transfected with 100nM siRNA diluted in a serum-free media with lipofectamine 2000 reagent (Thermo Fischer) in a 6 well plate as per the manufacturer’s protocol. The transfected medium was replaced with complete medium (10% FBS and 1% penicillin/streptomycin) after 9 hrs of transfection. Cells were harvested for RNA isolation after 36 hrs of post-transfection, and the knockdown efficiency for Hmrhl was verified by real-time PCR. List of all RT-PCR primers, si-RNA and probes used for the experiments are listed in Table S1.

### Hmrhl stability assay

To test the Hmrhl stability, 3×10^5^ cells were cultured with complete media (10% FBS and 1% penicillin-streptomycin solution) in a 6-well plate for 24 hrs. Subsequently, the media was replaced by fresh medium containing 10ug/ml of Actinomycin D (Invitrogen) and was incubated for different time intervals at 37°C. RNA was isolated for each time point and real-time PCR was performed to measure the expression of Hmrhl along with 18s rRNA as control.

### Subcellular fractionation

Subcellular fractionation was performed by following the previously described protocol by Gagnon et al., 2014 (45). Briefly, 5-10 million K562 cells were lysed in 380μl of ice-cold HLB (Hypotonic lysis buffer: 10 mM Tris, pH 7.5, 10 mM NaCl, 3 mM MgCl_2_, 0.3% NP-40 and 10% glycerol supplemented with 100 U of Ribolock RNase inhibitor and incubated on ice for 10min followed by centrifugation at 1000g at 4°C for 3 min. The supernatant containing cytoplasmic fraction was kept separately, and the nuclear pellet was washed with 1 ml of ice-cold HLB buffer three times and centrifuged at 300 g at 4°C for 2 min. To isolate total nuclear and cytoplasmic RNA, 1 ml of TRIzol reagent (Thermo Fisher Scientific) was added directly to the fractions and proceeded for RNA extraction. To separate nucleoplasmic and chromatin fractions, the nuclear pellet was resuspended in 380μl of MWS buffer (Modified Wuarin-Schibler buffer: 10 mM Tris-HCl, pH 7.0, 4 mM EDTA, 0.3 M NaCl, 1 M urea, and 1% NP-40), supplemented with 100 U of Ribolock RNase inhibitor and incubated on ice for 10 min followed by centrifugation at 1000g for 3 min at 4°C. The supernatant was kept as the nucleoplasmic fraction and the chromatin pellet was washed with ice-cold MWS buffer, centrifuged at 500g for 2 min at 4°C. TRIzol was then added to both nucleoplasmic fraction and chromatin pellet and proceeded for RNA extraction, as described earlier.

### Stellaris RNA fluorescent in situ hybridization (FISH)

RNA-FISH was carried out by following the protocol from Biosearch Technologies (https://www.biosearchtech.com/products/rna-fish). A set of Stellaris RNA FISH Probes labeled with Cy5 dye specific to Hmrhl targeted transcript was purchased from Biosearch Technologies. Briefly, 5×10^6^ cells were washed with PBS, followed by fixation in 1 ml fixation buffer (37% formaldehyde in PBS) for 10 min at room temperature. Cells were then centrifuged and washed thrice with PBS before permeabilizing with 70% ethanol for at least 1 hr at 2-8°C. Cell pellet was washed once with washed buffer A (Biosearch Technologies, Cat# SMF-WA1-60) and then proceeded for hybridization with 100ul hybridization buffer (Biosearch Technologies, Cat# SMFHB1-10) containing Hmrhl probe (working concentration 125 nM) for overnight at 37°C. Next day, cells were washed once with wash buffer A and resuspended in the same to be incubated for 30 min at 37°C in the dark. After incubation, cells were centrifuged and resuspended in wash buffer A containing DAPI stain and incubated further for 30 mins at 37°C in the dark. Next, cells were washed with wash buffer B (Biosearch Technologies, Cat# SMF-WB1-20) and resuspend in a small drop (approximately 30 µl) of Vectashield Mounting Medium. Subsequently, 10 ul of cell suspension was mounted on a slide for microscopic observation.

### Cell proliferation assay

Cell proliferation was evaluated using the Cell Counting Kit-8 (CCK8) assay (Dojindo). Cells (5000 per 100μl) with or without Hmrhl silencing were used, and cell proliferation was detected at every 24 hrs up to 96 hrs according to the manufacturer’s protocol. Briefly, 10μl of CCK 8 solution was added to each sample (5000cells/100μl) and was incubated for 2 hrs at 37°C. The colour developed was then measured spectrophotometrically (VERSAmax microplate reader) at 450 nm. Each sample was taken in triplicates, and the experiment was repeated with three biological replicates. After subtraction of background, the cell proliferation was represented as net absorbance.

### Transwell Assays

#### Migration assay

The migration of K562 cells was assessed using 8μm inserts (Corning Incorporated, Corning, NY, USA) according to the manufacturer’s instructions. Cells (1.5×10^5^) with/without silenced Hmrhl were suspended in the 500 μl of serum-free RPMI 1640 media and added onto the upper chambers of the inserts. Then, the inserts were placed in a 24-well plate, and the lower chambers were filled with 750 μl of RPMI 1640 media containing 20% FBS, which served as the chemoattractant. Cells were then allowed to migrate for 24 hrs to 48 hrs. Thereafter, the migrated cells in the bottom chamber were stained with 0.4 % Trypan Blue (Sigma) for 1 min, counted in a hemocytometer under an inverted microscope (at 10X in Olympus CKX41), and expressed as a percentage of cells migrated as compared with control. Also, migrated cells were quantified using CCK-8 assay as described above, and the results were represented as percentage cell migration for each group compared to control.

#### Invasion Assay

The invasiveness of K562 cells was monitored using invasion chambers (24well) coated with BD Matrigel™ matrix (8μm pore size, Corning). The invasion was measured by determining the ability of K562 cells to migrate through the Matrigel (a reconstituted basement membrane). Briefly, the matrigel membrane was first rehydrated by adding 500μl of warm (37°C) media (RMPI-1640 with 10% FBS) in both the chambers of the insert and incubating it for 2 hrs at 37°C. After rehydration 750 μL RPMI-1640 with 20% FBS, was added to the lower compartment of the invasion chamber while K562 cells (2.5×10^5^, with/without silenced Hmrhl) were suspended in 500μL serum-free RPMI-1640 and were seeded in the upper compartment of the invasion chambers and incubated for 48 hrs. After 48 hrs, cells that invaded to the bottom chamber were analyzed by manual cell counting using 0.4% trypan blue as well as by CCK8 assay. Results are represented as the percentage of invaded cells in different groups as compared to the control sample.

### Cell apoptosis assay

For analysis of cellular apoptosis, instructions provided by the kit (Annexin V-FITC Apoptosis Detection Kit, Sigma, Catalog No. APOAF-50TST) was followed. Cells (with/without Hmrhl silencing) were collected, rinsed twice with cold PBS, and then resuspended in 1X Binding Buffer at a concentration of 1×10^6^ cells/ml. Cells were then stained with annexin-FITC and PI and incubation at RT (25°C) for 15 min in the dark. FACS analysis was performed using flow cytometer (BD FACS Calibur). 10,000 reads were taken in the flow cytometer. For positive control of apoptosis, cells were treated with Imatinib (1µM) for 48 hrs. Cells that stained positive for annexin-FITC were either in the end stage of apoptosis or undergoing necrosis.

### Cell cycle analysis

K562 cells (1×10^6^ cells/ml) from different experimental groups were collected and washed twice with PBS. Subsequently, the cells were fixed in 70% ethanol for 30 min. After washing again twice with PBS, cells were finally resuspended in PBS and treated with 2 µl RNase A (10mg/ml, boiled for 5 min, aliquoted and stored frozen at −20°C) for 2 hrs at 37°C. 10 µl of Propidium Iodide (50 μg/ml, BD Biosciences) was added to the mixture. Flow cytometry was performed to detect and analyze changes in the cell cycle distribution using a flow cytometer (BD FACSCalibur).

### Chromatin immunoprecipitation (ChIP)

For the assay cells (1×10^6^) were fixed with 1% formaldehyde for 10 min at RT and the crosslinking was subsequently quenched with 125 mM glycine for 5 min at RT. After washing with PBS containing EDTA-free Complete Protease Inhibitor Cocktail (Roche) twice, cell pellets were resuspended in 300 µl of lysis buffer (50 mM Tris, pH 8.0; 10 mM EDTA; 1% SDS containing Protease Inhibitor Cocktail) and were allowed to sit on ice for 10 min. Cell extracts were then sonicated at high intensity for 30 min with 30 sec on/off cycles in a Bioruptor sonicator (Diagenode). Sonicated samples were diluted (1:10) in ChIP dilution buffer (1.1% Triton X-100; 1.2 mM EDTA; 16.7 mM Tris, pH 8.0; 167 mM NaCl, Protease Inhibitor Cocktail) and centrifuged at 20,000g for 10 min at 4°C. An aliquot (50 µl) of the soluble chromatin was kept as input. The chromatin was precleared with beads and immunoprecipitated with Tal1antibody (Diagenode, cat no: C15200012) overnight at 4°C. Next day 40 µl of protein A dynabeads (Invitrogen) were added to the immunocomplex and incubated at 4°C for 2 hrs. Beads were washed with Low salt buffer (150 mM NaCl; 0.1% SDS; 1% Triton X-100; 2 mM EDTA; 20 mM Tris, pH 8.0), high salt buffer (500 mM NaCl; 0.1% SDS; 1% Triton X-100; 2 mM EDTA; 20 mM Tris, pH 8.0), lithium chloride buffer (0.25 M LiCl; 1% NP-40; 1% sodium deoxycholate; 1 mM EDTA; 10 mM Tris, pH 8.0) and TE buffer (10 mM Tris, pH 8.0; 1 mM EDTA). The immunocomplex was eluted in elution buffer (1% SDS and 0.1 M NaHCO3) and reverse crosslinked overnight at 65°C followed by proteinase K digestion at 45°C for 1 hr. Subsequently the DNA was extracted and subjected to real-time PCR.

### Chromatin Isolation by RNA Purification (ChIRP)

ChIRP was carried out according to the previously described protocol by Chu et al., 2012 (46). Hmrhl specific anti-sense probes with BiotinTEG at 3’ end was designed using the Biosearch Technologies ChIRP Probe Designer tool (https://www.biosearchtech.com/chirpdesigner/). As a negative control for ChIRP, the LacZ probe with BiotinTEG at its 3’ end was used. About 60 million cells were harvested for a single ChIRP reaction. In brief, cells were cross-linked with 1% glutaraldehyde for 10 min at room temperature. Crosslinking cells were then quenched with 0.125 M glycine for 5 min, followed by centrifugation at 2000g to collect the cell pellet. Cells were then lysed using freshly prepared lysis buffer (50 mM TrisCl, pH 7.0, 10 mM EDTA, 1 % SDS) supplemented with 1mM PMSF, 1X protease inhibitor cocktail (PI) and Superase. The suspension was then sonicated in a Biorupture sonicator with 30 seconds ON and 30 seconds OFF cycle for 60 minutes. Sonicated chromatin was then centrifuged at 16000g for 10 minutes at 4°C. For a typical ChIRP reaction, 1 ml of sonicated lysate was used, in which 5% was kept aside as RNA and DNA input. Sonicated chromatin was hybridized with hybridization buffer (750 mM NaCl, 1 % SDS, 50 mM Tris-Cl pH 7.0, 1 mM EDTA, 15 % formamide) supplemented with 1mM PMSF, 1X PI and Superase. Hmrhl probe (100 pmol), as well as LacZ probe was added per 1ml of chromatin lysate in two separate tubes and was incubated for 4 hrs at 37°C with constant mixing. Approximately, 100 μl C-1 magnetic beads were then added to the 1ml chromatin lysate and incubated further for 30 min at 37°C with constant mixing. The beads were then washed five times with wash buffer (2xSSC, 0.5 % SDS, 1 mM PMSF). From the last wash, 100 μl was set aside for RNA isolation and 900 μl for DNA isolation. Hmrhl pull-down efficiency was verified by qRT-PCR analysis. For ChIRP sequencing, DNA was isolated according to the said protocol and sent for high throughput sequencing in duplicates.

### RNA-Seq analysis

GRCh38 (hg38) Genome was downloaded from GENCODE and indexed using Bowtie2-build with default parameters. Adapter ligation was done using Trim Galore (v0.4.4) and each of the raw Fastq files were passed through a quality check using the Fast QC. PCR duplicates were removed using the Samtools 1.3.1 with the help of rmdup option. Each of the raw files was then aligned to GRCh38 (hg38) genome assembly using TopHat with default parameters for paired end sequencing as described in Trapnell et al., 2012 (47). After aligning, quantification of transcripts was performed using Cufflinks and then Cuffmerge was used to create merged transcriptome annotation. Finally, differentially expressed (DE) genes were identified using Cuffdiff. The threshold for DE genes was log_2_ (fold change) >1.5 for up-regulated genes and log_2_ (fold change) <1.5 for down-regulated genes. GO enrichment analysis Gene Ontology (GO) analysis was performed in PANTHER (48). Significant enrichment test was performed with the set of differentially expressed genes in PANTHER.

### Cluster analysis

The Hierarchical clustering method was performed using Cluster 3.0 (49). Gene expression data (FPKM of all samples i.e, scrambled and siRNA treated) was taken and log_2_ transformed. Low expressed (FPKM<0.05) and invariant genes were removed. Then genes were centered, and clustering was performed based on the differential expression pattern of genes and fold change. Genes were grouped in 11 clusters and visualized as a network in Cytoscape (50). The Functional enrichment of each cluster was performed using the Gene Mania Tool (51). Heatmaps were generated in Java TreeView 3.0 for both up and down-regulated genes.

### TF network analysis

Among all DE protein-coding genes, the transcription factors were identified using PANTHER protein class module. Motifs were downloaded for all transcription factors from JASPAR (52) and sequence of interest for each TF (1.5 kb upstream & 500bp downstream of TSS) was extracted using BedtoFasta of the Bedtools suite (53). Then each motif was scanned across the sequence of all TFs to create the table matrix that reflects the number of binding sites for each TF across the other TFs using MEME suite (54) with an e-value of 1E-04. Finally, the heatmap was generated from the table matrix using R 3.3.2.

### ChIRP-Seq analysis

GRCh38 (hg38) Genome was downloaded from GENCODE and indexed using Bowtie2-build with default parameters. Adapter ligation was done using Trim Galore (v 0.4.4) and each of the raw Fastq files were passed through a quality check using the FastQC. PCR duplicates were removed using the Samtools 1.3.1 with the help of ‘rmdup’ option. Each of the raw files was then aligned to GRCh38 (hg38) genome assembly using Bowtie 2 with default parameters for paired-end sequencing as described in Trapnell et al., 2012 (47). Replicates of both control and treated were merged respectively using the Samtools 1.3.1. Peaks were called using MACS2. Final peaks were selected, giving the criteria of above 5-fold change and p value < 0.05. Hmrhl Motif Prediction Motifs were identified using MEME, based on the criteria of One Occurrence Per Sequence (OOPS) and significance of 1E -04 for all ChIRP-Seq genomic loci. Sequence for each locus was extracted from GRCh38 (hg38) genome using bedtofasta of bedtools suite. After feeding sequences from the genomic loci obtained from MACS2, two significant motifs were identified in MEME motif discovery suite.

### Triplex Prediction

Sequences from the Hmrhl occupied regions (in addition extended upto +/− 25 bp) of the selected genes (TP53, PDGFRB, ZIC1) were used to assess the compatibility of potential TFO/TTS pairs according to the canonical triplex formation rules between the entire stretch of Hmhrl sequence (Triplex Forming Oligo Nucleotide) and the aforementioned Hmrhl occupied regions of a respective genes (Triplex Target Sites) using the software Triplexator with default parameters (55).

## RESULTS

### Expression of LncRNA Hmrhl is co-related with chronic myelogenous leukemia (CML)

In our recently published study on Hmrhl, we had observed that most significant upregulation in the expression of Hmrhl was in lymphocyte tumor samples among all the other cancers (44). We went ahead and looked for the expression pattern of Hmrhl in patient samples of the two types of leukemia namely chronic myelogenous leukaemia (CML, slow developing) and acute myeloid leukaemia (AML, fast developing and more severe). For this purpose we took advantage of the publicly available transcriptome data of both CML and AML patients (from https://www.ebi.ac.uk/ega/dacs/EGAC00001000481) and calculated the corresponding transcript FPKM values. Based on our data analysis, Hmrhl expression was indeed found to be upregulated in CML patient samples in contrast to the samples from AML patients (Figure 1A). We had previously shown that Hmrhl expression is elevated in K562 cells (derived from myelogenous leukemia cell line) in comparison with the control cell line GM12878. We have reconfirmed this observation as shown in Figure 1B. Thus, we have selected K562 cell line to delineate potential molecular function of Hmrhl in CML in the present study.

**Figure 1:**
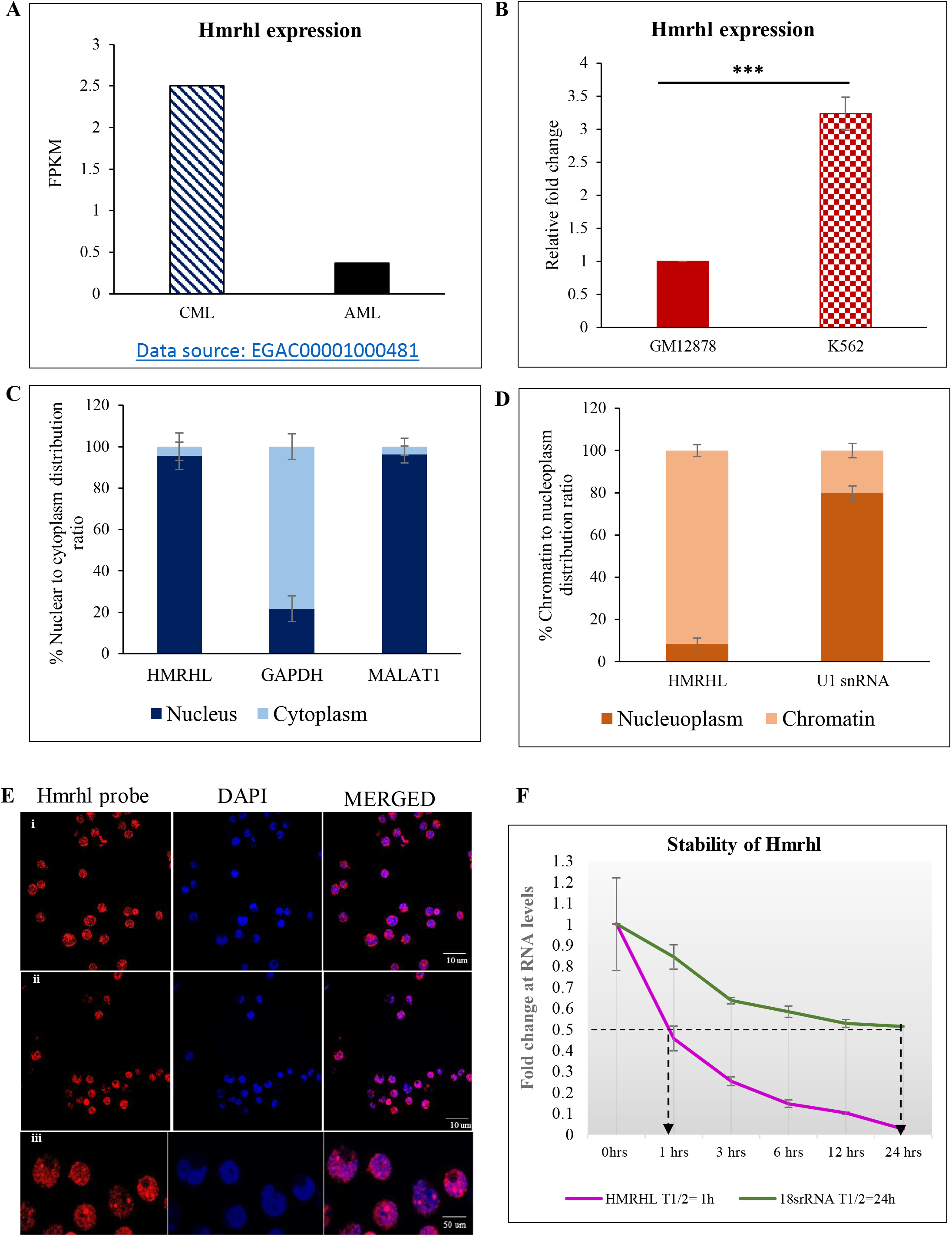
General characterization of LncRNA Hmrhl. (A) FPKM values depicting level of lncRNA Hmrhl in AML and CML patient samples as obtained from public data source “https://www.ebi.ac.uk/ega/dacs/EGAC00001000481”. (B) RT-qPCR analysis of Hmrhl RNA levels with respect to internal control GAPDH in CML cell line K562 and control cell line GM12878. (C) Subcellular fractionation showing nuclear restricted expression of Hmrhl RNA with levels of GAPDH and MALAT1 serving as authentication of purity for each fraction. (D) Chromatin association of Hmrhl RNA as represented by sub nuclear distribution of Hmrhl compared to nucleoplasm associated U1snRNA. (E) RNA-FISH using lncRNA Hmrhl specific probe reconfirms nuclear restriction of Hmrhl in K562 cells. (F) RNA stability assay of Hmrhl RNA and 18S rRNA in the presence of actinomycin D (10ug/ml). The data are presented as mean ± SD from three independent experiments, *p<0.05, **p<0.01, ***p<0.001 by student’s t-test.

### Hmrhl is a chromatin bounded LncRNA with a short half-life

Since, we had shown earlier that the mouse mrhl RNA is nuclear restricted and has regulatory function at chromatin level, we were curious to examine the sub cellular localization of Hmrhl RNA in K562 cells (43). Results from biochemical fractionation studies indicate the presence of Hmrhl predominantly in the nuclear fraction which was in parallel with MALAT1, a well-known highly abundant nuclear enriched lncRNA in human (Figure 1C) (31). GAPDH was used as a negative control to authenticate the purity of the nuclear fraction (Figure 1C). Hmrhl was further found to be predominately associated with the chromatin fraction with very little in the nucleoplasm fraction (Figure 1D). U1snRNA served as a positive control for nucleoplasm in the subnuclear fractionation study (Figure 1D). RNA FISH experiment further validated Hmrhl nuclear localization and it appears as bright red punctuate foci on blue DAPI background of chromatin mass (Figure 1E). We also observed some spots to be of brighter intensity as compared to other foci suggesting that Hmrhl might be highly concentrated at some of the genomic loci (Figure 1E). This corroborated our earlier study where Hmrhl was also found to be nuclear restricted and chromatin associated in HEK-293T cells (44).

The stability of noncoding RNAs (ncRNAs) like protein-coding mRNAs, is found to be closely associated with their physiological function, in general regulatory RNAs been less stable than the one with housekeeping function (56). In addition, global measurement of the half-lives of lncRNAs in mouse has revealed that spliced lncRNAs are more stable than unspliced lncRNAs (single exon), cytoplasmic lncRNAs are more stable than nuclear and antisense-overlapping lncRNAs are more stable than those transcribed from the intron (57). Since Hmrhl is an unspliced, nuclear, intronic lncRNA, we presumed Hmrhl to be a short lived lncRNA. To find out the exact half-life of Hmrhl, we used actinomycin D method and evaluated the half-life against highly stable 18S rRNA as positive control (Figure 1F). Results shown in Figure 1F indicate that Hmrhl certainly is a short lived lncRNA with a half-life of approximately 1 hr.

### Perturbation of Hmrhl affects the genes involved in key signaling pathway in leukemia as well as in the developmental process

To elucidate the functional role of Hmrhl, we took the RNA-seq approach to examine whether it regulates the expression of protein coding genes in these cells. Hmrhl was silenced using a pool of 4 targeted siRNA in K562 cells which gave us an average down regulation efficiency of approximately 60% as compared to the scrambled control siRNA (Figure 2A and B). Both Hmrhl siRNA and scramble siRNA treated cells were then subjected to RNA-seq analysis. Our analysis with a fold cut-off of 1.5 +/− revealed a total of 1584 dysregulated genes (DE) up on Hmrhl silencing, out of which 831 genes were found to be protein coding (Figure 2C, Supplementary file 1). Further evaluation of protein coding DE genes indicated that majority of them are down regulated (592) compared to upregulated genes (239) (Figure 2C).

**Figure 2:**
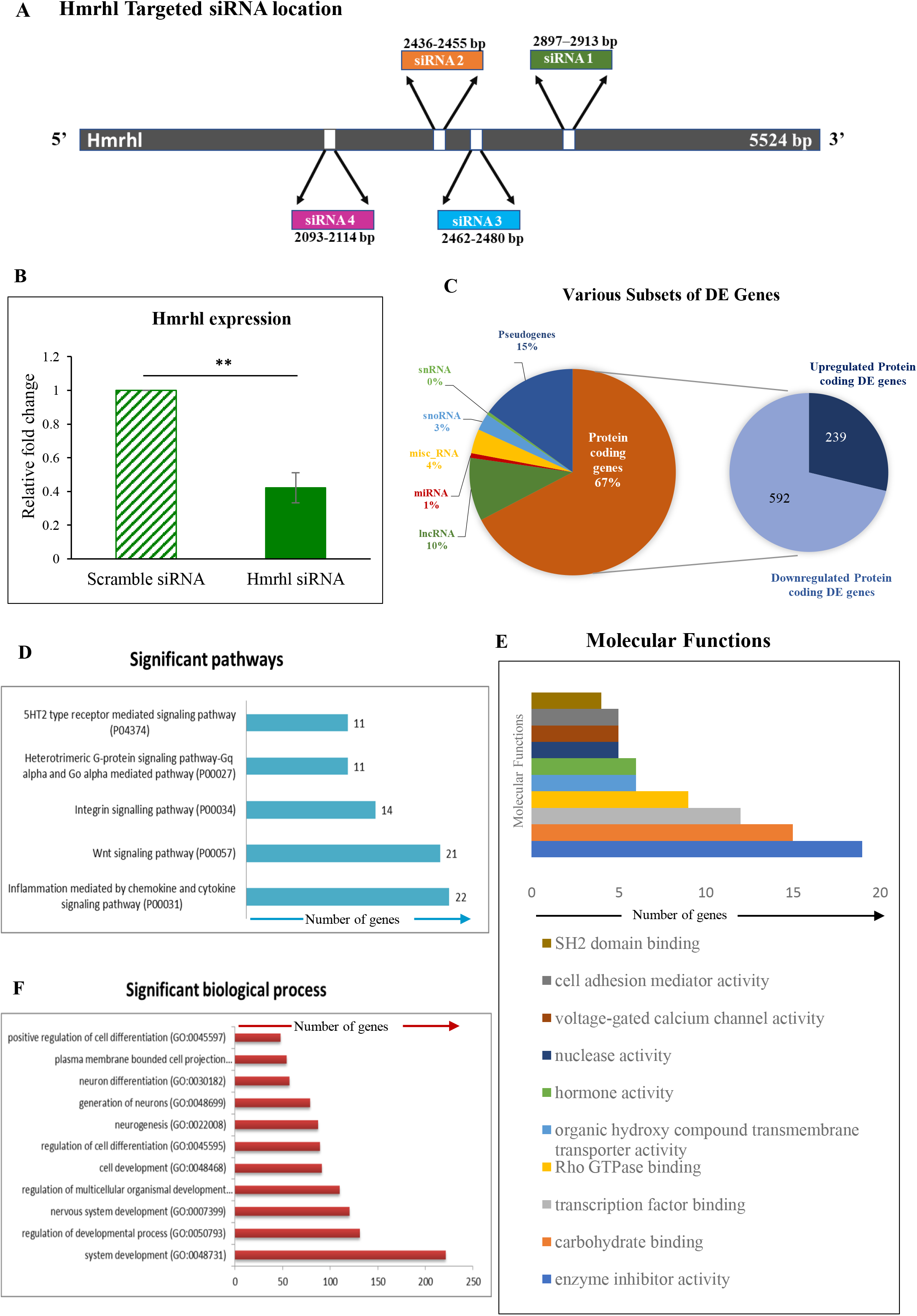
Transcriptome analysis reveals role of lncRNA Hmrhl in biological processes related to development and signaling. (A) Diagrammatic representation of targeted siRNA binding sites used for silencing Hmrhl. (B) Knockdown efficiency of Hmrhl shown as relative expression of Hmrhl after si-Hmrhl and scramble-RNA treated K562 cells with GAPDH as internal control. (C) Diagrammatic representation of percentage of DE genes in each category obtained after RNA-seq analysis of transcriptome after perturbation of Hmrhl RNA. (D), (E) and (F) Gene ontology (GO) enrichment analysis of DE genes showing number of genes associated with significant pathways, molecular functions and key biological processes respectively. Error bars indicate standard deviation from three independent experiments. *p<0.05, **p<0.01, ***p<0.001 by student’s t-test.

Following the identification of differentially expressed genes, Gene Ontology (GO) enrichment analysis of protein coding genes with a p-value <0.05 was carried out to identify pathways that are perturbed upon Hmrhl silencing. Notably, several pathways that are associated with key signaling processes in cancer were found to be affected such as chemokine and cytokine signaling pathway (P00031, 22 genes) followed by the Wnt signaling pathway (P00057, 21 genes) and integrin signaling pathway (P00034, 14 genes) which were found be most enriched along with heterotrimetric G-protein signaling pathway (P00027, 11 genes) and 5HT2 type receptor mediated signaling pathway (P04374, 11 genes) in the top five list (Figure 2D, Table 1). Moreover, GO analysis also revealed diverse molecular functions and a number of biological processes to be altered upon Hmrhl silencing. Figure 2E shows top 10 of such enriched molecular functions which includes enzyme inhibitor activity (19 genes, p-val:0.0154), transcription factor binding (12 genes, p-val:0.0136), Rho GTPase binding (9 genes, p-val:0.0329), nuclease activity (5 genes, p-val:0.00225), voltage-gated calcium channel activity (5 genes, p-val:0.0214) and cell adhesion mediator activity (5 genes, p-val:0.0374). We also observed from the analysis that most of the genes with altered expression pattern belongs to biological processes of cellular development and differentiation such as system development (GO:0048731), developmental regulation process (GO:0050793), nervous system developmental process (GO:0007399), regulation of cell differentiation (GO:0045595) (Figure 2F). We could conclude, from our observations therefore, that Hmrhl is mostly involved in key signaling pathways and processes of cell development and differentiation in K562 cells.

**TABLE 1:** Detail description of DE genes in biological pathways as obtained after GO analysis

| **Inflammation mediated by cytokines and chemokines signaling pathway** |  |  |
| --- | --- | --- |
| **Gene** | **Fold change** | **Biological function** |
| RGS1 | 1.57636 | Regulates G protein-coupled receptor signalling cascades |
| CCL3 | -1.71239 | Monokine with inflammatory and chemokinetic properties. |
| ITGB2 | -1.22457 | Integral cell-surface proteins that participate in cell adhesion as well as cell-surface mediated signalling |
| IL6 | 1.57968 | Cytokine with a wide variety of biological functions. |
| GNA15 | -5.38507 | Involved as modulators or transducers in various transmembrane signalling systems |
| CCR4 | 3.77073 | Function as a chemoattractant homing receptor |
| FPR2 | -1.78241 | Powerful neutrophil chemotactic factors |
| MYH7B | -2.16246 | Involved in muscle contraction. |
| PRKCE | -2.62241 | Known to be involved in diverse cellular signalling pathways |
| MYH11 | -2.30486 | Involved in muscle contraction. |
| CXCL10 | -2.27738 | Involved in a wide variety of processes such as chemotaxis, differentiation, and activation of peripheral immune cells, regulation of cell growth, apoptosis |
| IL1B | -3.97255 | An important mediator of the inflammatory response, and is involved in a variety of cellular activities, including cell proliferation, differentiation, and apoptosis. |
| ITGA2 | -1.27507 | It is responsible for adhesion of platelets and other cells to collagens |
| GNA14 | -1.21527 | Involved as modulators or transducers in various transmembrane signalling systems |
| GNG7 | -3.67311 |  |
| GNB3 | -1.60354 |  |
| FPR1 | -1.25003 | Powerful neutrophil chemotactic factors |
| RELB | -1.27448 | Involved in many biological processes such as inflammation, immunity, differentiation, cell growth, tumorigenesis and apoptosis. |
| PLCB4 | 2.48726 | Important role in the intracellular transduction of many extracellular signals in the retina |
| PTAFR | -2.05699 | A chemotactic phospholipid mediator that possesses potent inflammatory, smooth-muscle contractile and hypotensive activity. |
| MYLK2 | -2.0379 | Implicated in the level of global muscle contraction and cardiac function |
| MYH3 | 2.22508 | Involved in cell movement and transport of materials within and between cells. |
| **WNT signaling pathway** |  |  |
| **Gene** | **Fold change** | **Biological function** |
| DCHS1 | -1.27603 | Calcium-dependent cell-adhesion protein. Mediates functions in neuroprogenitor cell proliferation and differentiation. |
| WNT8B | -1.68662 | May play an important role in the development and differentiation of certain forebrain structures, notably the hippocampus |
| PCDH12 | 2.29283 | Acts as a regulator of cell migration, probably via increasing cell-cell adhesion |
| GNA15 | -5.38507 | Involved as modulators or transducers in various transmembrane signalling systems. |
| MYH7B | -2.16246 | Involved in muscle contraction. |
| PRKCE | -2.62241 | Plays essential roles in cancer cell invasion and regulation of apoptosis, regulation of multiple cellular processes such as cell adhesion, motility, migration and cell cycle |
| ANKRD6 | 3.17182 | Allows efficient phosphorylation of beta-catenin, thereby inhibiting beta-catenin/Tcf signals. |
| CTNNA2 | 1.67243 | Regulate cell-cell adhesion and differentiation in the nervous system |
| CTNNA3 | -2.54244 | May be involved in formation of stretch-resistant cell-cell adhesion complexes. |
| GNA14 | -1.21527 | Involved as modulators or transducers in various transmembrane signalling systems. |
| CDH12 | -2.78544 | Calcium-dependent cell adhesion proteins with a role during a critical period of neuronal development |
| GNG7 | -3.67311 | Involved as a modulator or transducer in various transmembrane signalling systems. |
| GNB3 | -1.60354 | Involved as a modulator or transducer in various transmembrane signalling systems |
| FZD9 | -1.53141 | Involved in beta-catenin canonical signalling pathway |
| TP53 | -1.89693 | Acts as a tumour suppressor in many tumour types; induces growth arrest or apoptosis depending on the physiological circumstances and cell type. |
| FAT2 | -1.23901 | Involved in the regulation of cell migration and proliferation |
| PLCB4 | 2.48726 | Plays an important role in the intracellular transduction of many extracellular signals in the retina |
| PCDH11Y | 1.70719 | Play a role in cell-cell recognition during development of the central nervous system |
| FRZB | -1.71826 | Function as modulators of Wnt signalling through direct interaction with Wnts. |
| MYH3 | 2.22508 | Involved in cell movement and transport of materials within and between cells. |
| SFRP5 | -2.18751 | Have a role in regulating cell growth and differentiation in specific cell types. |
| **Interigin signaling pathway** |  |  |
| **Gene** | **Fold change** | **Biological function** |
| ITGB2 | -1.22457 | Involved in leukocyte adhesion and transmigration of leukocytes including T-cells and neutrophils |
| CAV1 | 2.14717 | scaffolding protein promotes cell cycle progression |
| COL9A2 | 2.03169 | extracellular matrix structural constituent conferring tensile strength |
| SH3D21 | -1.33154 | structural component of nucleus and plasma membrane |
| COL11A2 | 3.24925 | May play an important role in fibrillogenesis by controlling lateral growth of collagen II fibrils. |
| ITGA2 | -1.27507 | responsible for adhesion of platelets and other cells to collagens |
| COL17A1 | 2.04848 | May play a role in the integrity of hemidesmosome |
| COL4A3 | 1.66658 | major structural component of glomerular basement membranes |
| COL9A3 | -1.43053 | Structural component of hyaline cartilage and vitreous of the eye |
| ACTN2 | -1.3298 | anchor actin to a variety of intracellular structures. |
| ITGA10 | 2.5105 | Acts as a receptor for collagen. |
| LAMA2 | 2.37888 | mediate the attachment, migration and organization of cells into tissues during embryonic development by interacting with other extracellular matrix components |
| LAMC1 | -2.12482 | mediate the attachment, migration and organization of cells into tissues during embryonic development by interacting with other extracellular matrix components |
| ELMO1 | 1.7877 | Involved in cytoskeletal rearrangements required for phagocytosis of apoptotic cells and cell motility. |
| **Heterotrimeric G-protein signalling pathway** |  |  |
| **Gene** | **Fold change** | **Biological function** |
| CHRM4 | -1.26805 | Mediates various cellular responses, including inhibition of adenylate cyclase, breakdown of phosphoinositides and modulation of potassium channels through the action of G proteins |
| RGS1 | 1.57636 | Regulates G protein-coupled receptor signalling cascades |
| PLCB4 | 2.48726 | Plays an important role in the intracellular transduction of many extracellular signals in the retina. |
| PRKCE | -2.62241 | Plays essential roles in the regulation of multiple cellular processes linked to cytoskeletal proteins, such as cell adhesion, motility, migration and cell cycle |
| CACNA1A | 3.46911 | Involved in a variety of calcium-dependent processes, including muscle contraction, hormone or neurotransmitter release, gene expression, cell motility, cell division and cell death. |
| CACNA1E | -1.17684 | Involved in a variety of calcium-dependent processes, including muscle contraction, hormone or neurotransmitter release, gene expression, cell motility, cell division and cell death. |
| GNA15 | -5.38507 | Involved as modulators or transducers in various transmembrane signalling systems. |
| GNGT1 | 2.02369 |  |
| GNA14 | -1.21527 |  |
| GNG7 | -3.67311 |  |
| GNB3 | -1.60354 |  |

### Gene co-expression and TF interaction analysis reveals distinct gene networks associated with Hmrhl RNA

We observed subsets of important transcription factors (TF) and common cancer related genes to be dysregulated during the silencing process of Hmrhl (Figure 3A & B) implying probable role of Hmrhl in regulation of these key TFs and cancer associated genes. A thorough literature search revealed many of these TFs and cancer related DE genes do have significant biological role in leukemia, development and signaling pathways. In TFs group (Figure 3B) for example, *KLF12*, an important regulator of gene expression during vertebrate development has been reported to be upregulated in K562 cells (58), while hypermethylation of *TFAP2E*, which is considered as a tumor suppressor gene was found to be associated with K562 cell line and shortened survival of CML patients (59, 60). Others, like *KLF2*, *MAFA*, *STAT4*, *ASCL2*, *KLF4*, *BATF*, *TP63* were all linked to leukemia and lymphomas in various studies (61–69). Many TFs, like *TBX8*, *FEV*, *SP8* with their crucial role in development were also identified (70–72). Expressed only in fetal hematopoietic cells and essential for hematopoietic stem cell renewal, *FEV*, was reported to be expressed in infant leukemia samples with involvement in maintenance and propagation of leukemic stem cells (72). Beside theses, others TFs were found to be associated with various signaling pathways which directly or indirectly modulates the cell growth, proliferation and apoptosis. For example, *ASCL2*, which is hypermethylated in lymphocytic leukemias and *SOX15*, a tumor suppressor, both acts via Wnt signaling pathway to modulate cell proliferation and invasion (64, 73, 74). Transcription factors, *STAT4* and *IRF5* both were reported to be expressed abnormally low in leukemia (63, 75). *STAT4*, which has association with neoplasia, is part of STAT signaling pathway critical for mediating the response of hematopoietic cells to a diverse spectrum of cytokines (76) while *IRF5*, which modulates expression of genes controlling cell growth and apoptosis, was recently demonstrated to be a target of BCR-ABL kinase activity which reduces CML cell proliferation (77). Another transcription factor KLF4 was reported to be critically regulating human myeloid leukemia cell proliferation and its dysregulated expression alters cell division, differentiation and apoptosis in myeloid leukemias (65, 66, 78). TP63, which regulates cellular activities like division, adhesion and apoptosis, was established to have a pro-survival effect with a role in blast crisis in CML (68, 69). This detailed analysis of DE TFs further supports our GO outcomes of RNA seq analysis where most effected pathways and biological processes belongs to development and signaling (Figure 2D & F). This was found true for the DE genes associated with cancer as well (Figure 3A). For example, both *GLI1* and *PTCH1* were found to be associated with hedgehog signaling, important for embryonic development and tumorigenesis (79, 80). Dysregulated expression and mutation in these genes has been corelated with phenotype and cancer stemness in CML (79, 80). CD274, also known as PDL1, is a receptor found on CML specific T-cells and are stipulated to increase immune suppressive signals leading to reduce antitumor activity within tumor microenvironment (81). It was also found to be associated with the phenotype and maintenance of leukemic stem cells in CML and AML (82, 83). Other DE genes, like *CCR4*, *ITK*, *LRP1B*, *GPC5*, *PDGRFβ*, *PRDM16*, *PTPRK*, *ZIC1* were all found to be involved in signaling pathways with fundamental role in development, immune response, cell growth, proliferation, migration etc. (84–91). Fusion transcripts, mutation and dysregulated expression of PDGRFB and PRDM16 were reported in many leukemias (89). Another study has reported the abnormal promoter methylation of PTPRK in 47% of ALL and B-cell lymphoma patients and was associated with decreased survival in the cohort (90).

**Figure 3:**
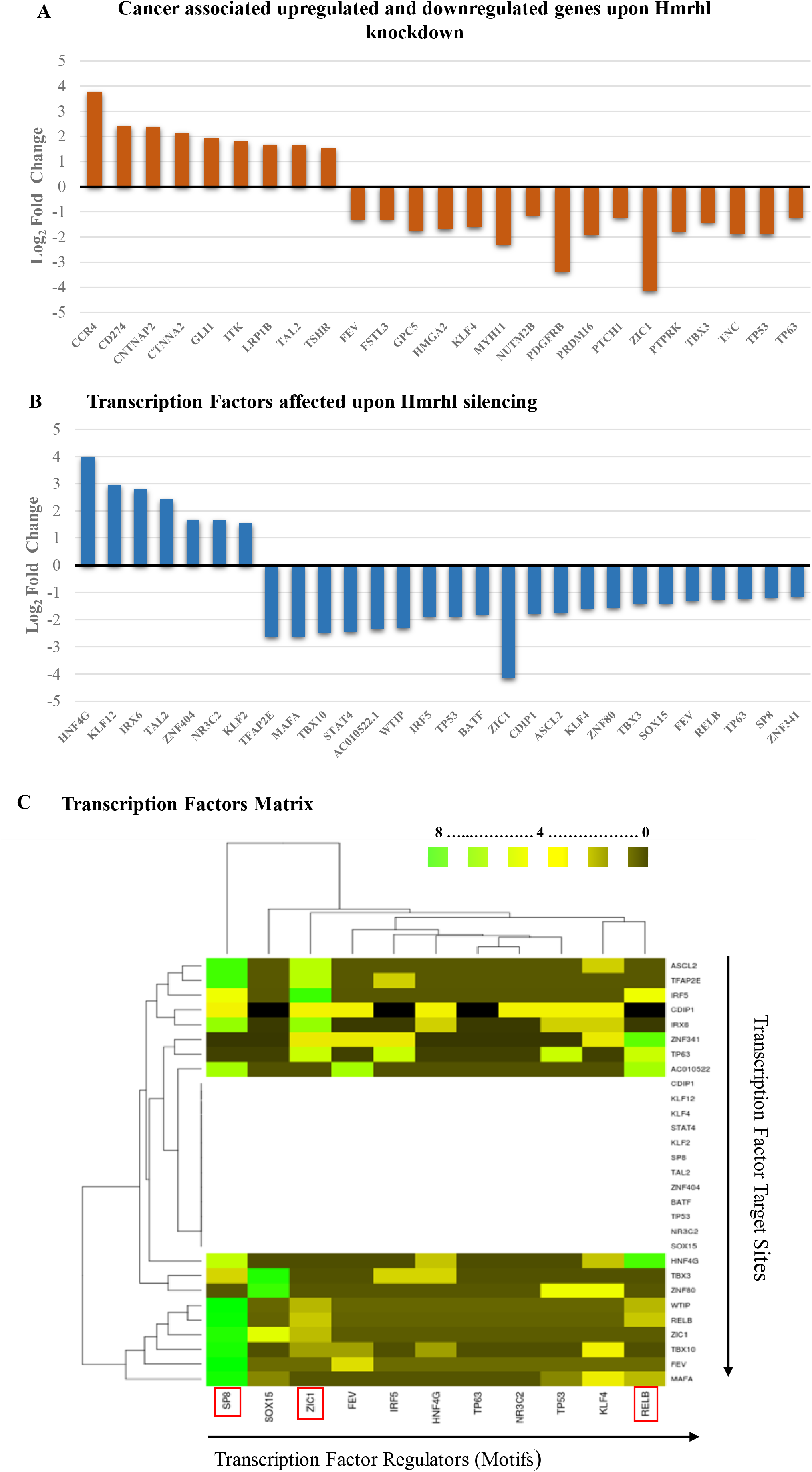
Further analysis of DE genes obtained after Hmrhl knockdown revealed some important (A) cancer associated genes and (B) Transcription factors. (C) Heat map visualization of DE Transcription factors matrix. Heat map was generated from the table matrix which was obtained using MEME suit.

We further carried out analysis of important TF regulators among all DE transcription factors obtained after Hmrhl silencing (Figure 3C, Supplementary file 2). Transcription factors (TFs) determine in large part the connectivity of gene regulatory networks as well as the quantitative level of gene expression. And it is evident from literature survey that dysregulation of TFs is associated with cancers (92). This allowed us to develop a TF matrix which shows SP8, ZIC1 and RELB as the top three dominating TFs (marked in red boxes, Figure 3C) as they have maximum number of motifs for binding across the promoters of all DE TFs (Table S2). Both, *SP8* and *ZIC1* genes have been shown to be essential for proper embryonic development, with *SP8* playing a key role in limb development whereas *ZIC1* is involved in neurogenesis (93–95). Dysregulation of both of these genes has been found to be associated with various cancers where they promote metastasis by regulating cell cycle and growth (91, 96). *RELB* on the other hand codes for a member of the nuclear factor kappa-B (NFKB) family of TFs and is involved in many biological processes such as inflammation, immunity, differentiation, cell growth, tumorigenesis and apoptosis (97, 98).

To understand the functional relevance of perturbed genes upon Hmrhl silencing in K562, we further screened DE genes to analyze gene-gene interactions to be identified as co-expression modules. We carried out the hierarchical clustering and obtained eleven clusters using the Cytoscape platform (Figure 4, Supplementary file 3). We subjected each of these gene cluster modules for functional enrichment with the Gene Mania tool, to obtain different cellular pathways to which these genes belong. Results from this analysis showed signal transduction and developmental activities for clusters 1; cluster 4 and 5 demonstrate ion channel and adhesion molecular activity. Nervous system functions for cluster 6 and kinase signaling processes for cluster 7 were obtained, whereas cluster 2, 9 and 10 were associated with activities for muscle contraction and potassium channel. Cluster 11 was found to be involved in the extracellular signaling process (Figure 4), whereas clusters 3 and 8 did not indicate any functional relevance to the cellular pathway (Figure S1).

**Figure 4:**
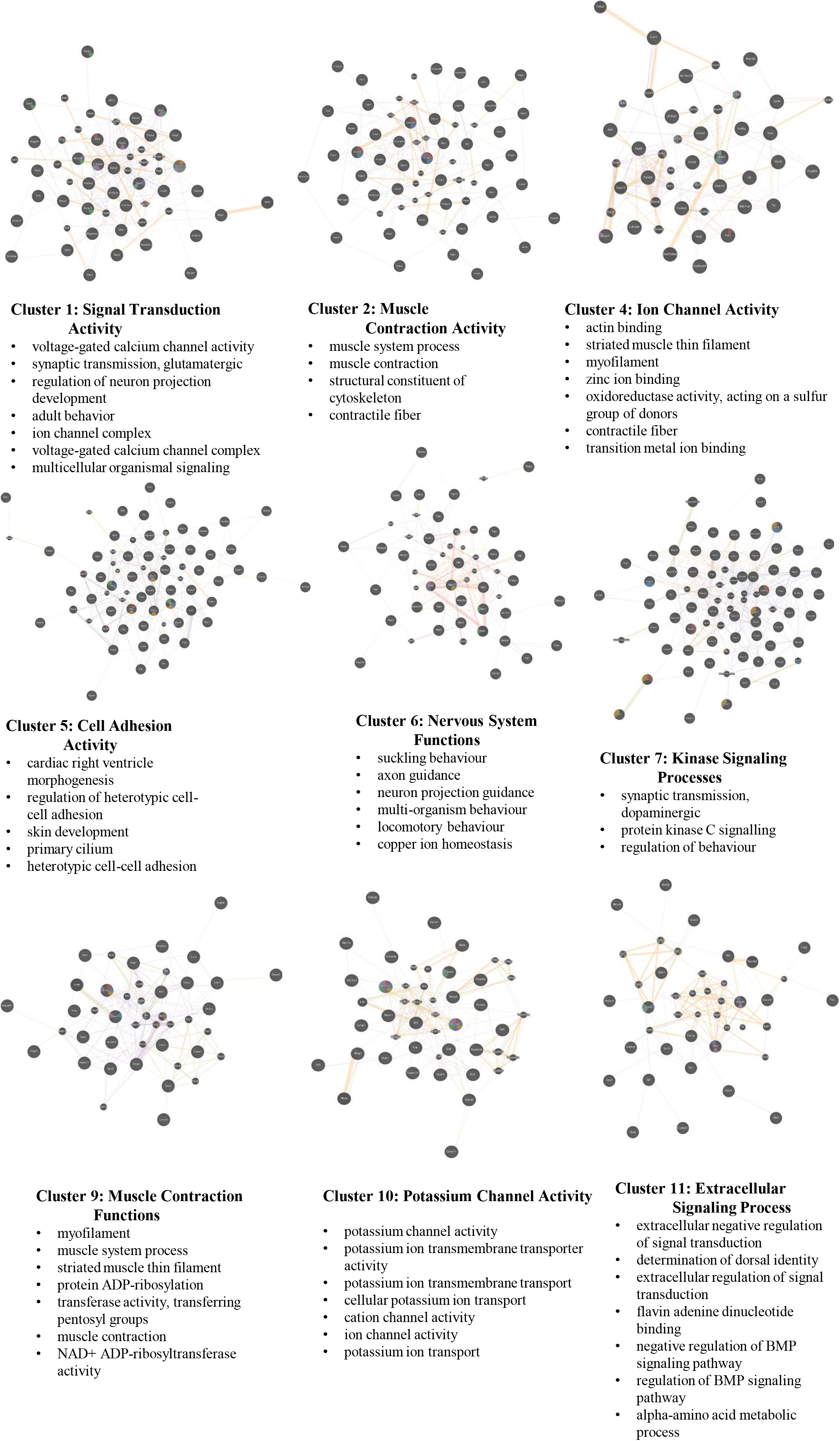
Gene co-expression modules (cluster 1, 2, 4, 5, 6, 7, 9, 10 and 11) showing their distinct functional profiles, obtained after hierarchical clustering using Cytoscape platform. ***See also Figure S1*:** Showing cluster 3 and 8 with no specific functional enrichment after gene co-expression analysis.

### Hmrhl RNA promotes cell proliferation, migration and invasion of K562 cells

We next investigated the cellular role of Hmrhl in cancer phenotype of K562 cells. Cell proliferation, migration and invasion are processes readily associated with cancer phenotype and studied widely to establish the role of a particular gene in cancer (99). Therefore, we started with CCK-8 assay which demonstrates significant suppression of K562 cell proliferation 48 hrs after Hmrhl silencing which continued till the time of observation i.e. 96 hrs (Figure 5A). We performed flow cytometric analysis to further answer the question whether down regulated Hmrhl condition affected proliferation of K562 cells is caused due to alteration in cellular apoptosis or cell cycle progression. We observed that down regulation of Hmrhl expression results in significant accumulation of cells at the G0/G1 phase (∼70%, p.val<0.01) as compared to control (∼50%) with reduction at S phase, from ∼30% in si-Hmrhl treated sample to ∼17% in control (p.val<0.001) and si-scramble (Figure 5B & C). However, no affect of Hmrhl silencing could be seen in apoptosis assay as all samples showed similar % of cells in each quadrant with maximum number of live cells and very few apoptotic cells (Figure S2A).

**Figure 5:**
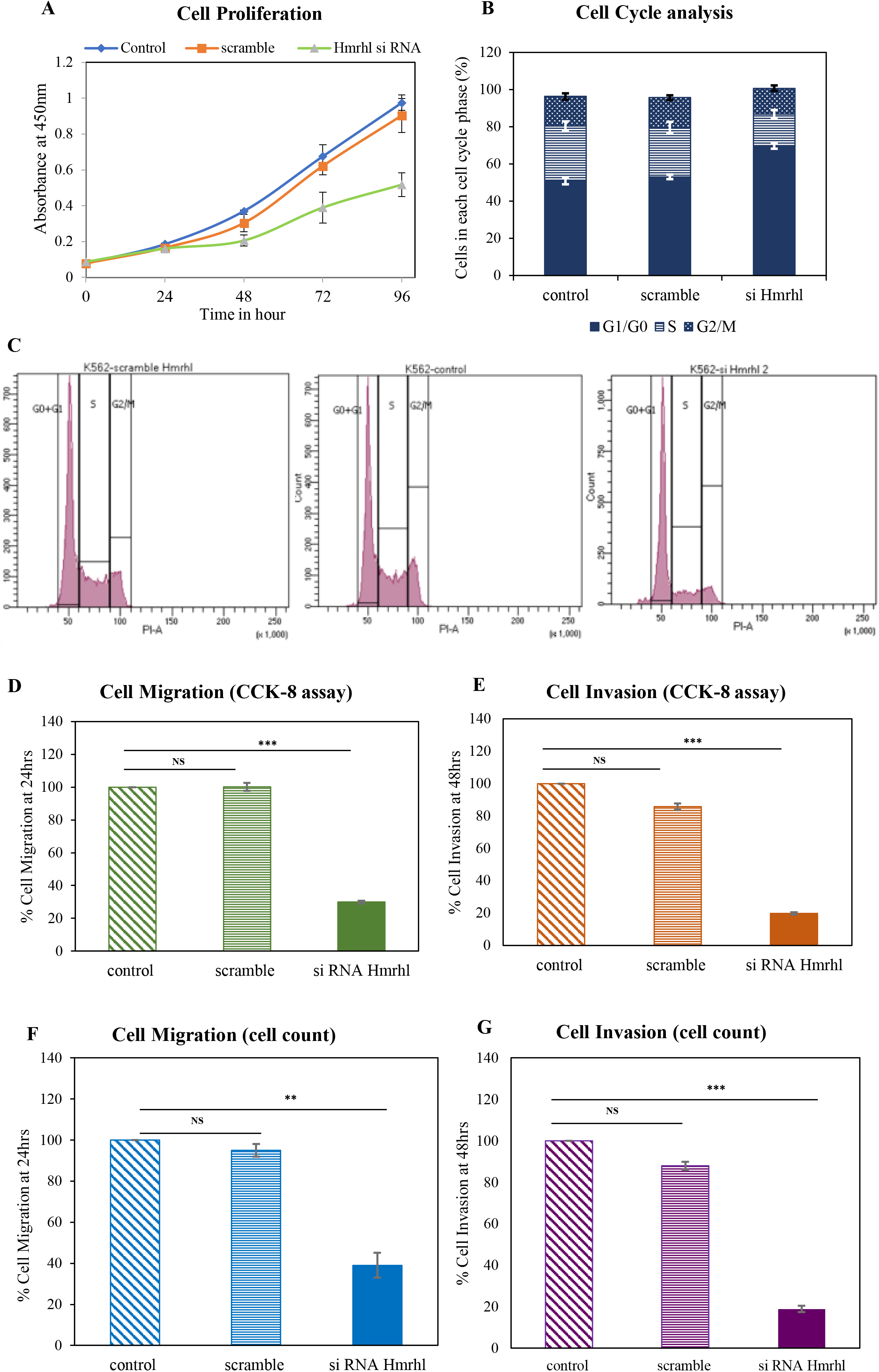
LncRNA Hmrhl contributes to cancer phenotype in K562. (A) CCK-8 assay shows decline in cell proliferation up on Hmrhl silencing measured over a time period of 96 hrs. (B) & (C) Bar graph and Flow cytometry histogram showing G0/G1 arrest represented as % of cells in each cell cycle phase after knockdown of Hmrhl compared to scramble RNA treatment and control samples. (D) & (E) Transwell assay followed by CCK-8 assay to evaluate the endpoint reveals inhibition of cellular migration and invasion after si-Hmrhl treatment in K562 cells respectively. (F) & (G) Reconfirmation of effect of Hmrhl knockdown on cell migration and invasion respectively using manual cell counting as the end point assay. The data are presented as mean ± SD from three independent experiments, *p<0.05, **p<0.01, ***p<0.001 by student’s t-test ***See also Figure S2*:** (A) Evaluation of cell death by apoptosis as an effect of Hmrhl silencing using annexin V-FITC/PI staining. Dot plot profile of (i) control cells (ii) si-Hmrhl treated K562 cells (iii) scramble-RNA treated cells. (B) & (C) Cell migration evaluated by CCK-8 assay and manual counting respectively shows no difference over the time period of 24 and 48hrs.

The results of the transwell assay showed that siRNA treatment significantly impaired the migration capability of treated cells as compared to control (Figure 5D). With a 70% inhibition in cell migration observed after 24 hrs of Hmrhl silencing, no further decline was noted till 48 hrs (Figure S2B). Hmrhl downregulated K562 cells also inhibited their invasion ability by approximately 80% when compared to control cells (Figure 5E). Both the assays were further verified using manual counting which showed similar results as obtained by CCK-8 after transwell assay (Figure 5F & G, Figure S2C).

### Hmrhl binding at chromatin level regulates differential expression of target gene via triplex formation

Since, Hmrhl is associated with chromatin, we were curious to identify the genomic binding sites by employing ChIRP-seq technique. Chromatin isolation by RNA purification (ChIRP) coupled with high-throughput sequencing (ChIRP-seq) is a robust technique that allows one to identifying genome-wide chromatin binding sites for any lncRNAs (46). We used biotinylated antisense oligonucleotides probes to pull down Hmrhl, specificity of which was confirmed by qRT-PCR against non-specific control probes against lacZ (Figure 6A). Sequencing of the ChIRP DNA indicated association of Hmrhl with thousands of genomic loci on chromatin. Highly significant sequence reads selected on the bases of 5-fold enrichment over input DNA and a p-value <0.01, gave us a total of 68,177 peaks (Supplementary file 4). Further annotation of these peaks revealed enrichment of Hmrhl to be significantly higher at some chromatin regions as compared to other loci, which includes intergenic, introns, repeat elements as well as promoter regions (Figure 6B).

**Figure 6:**
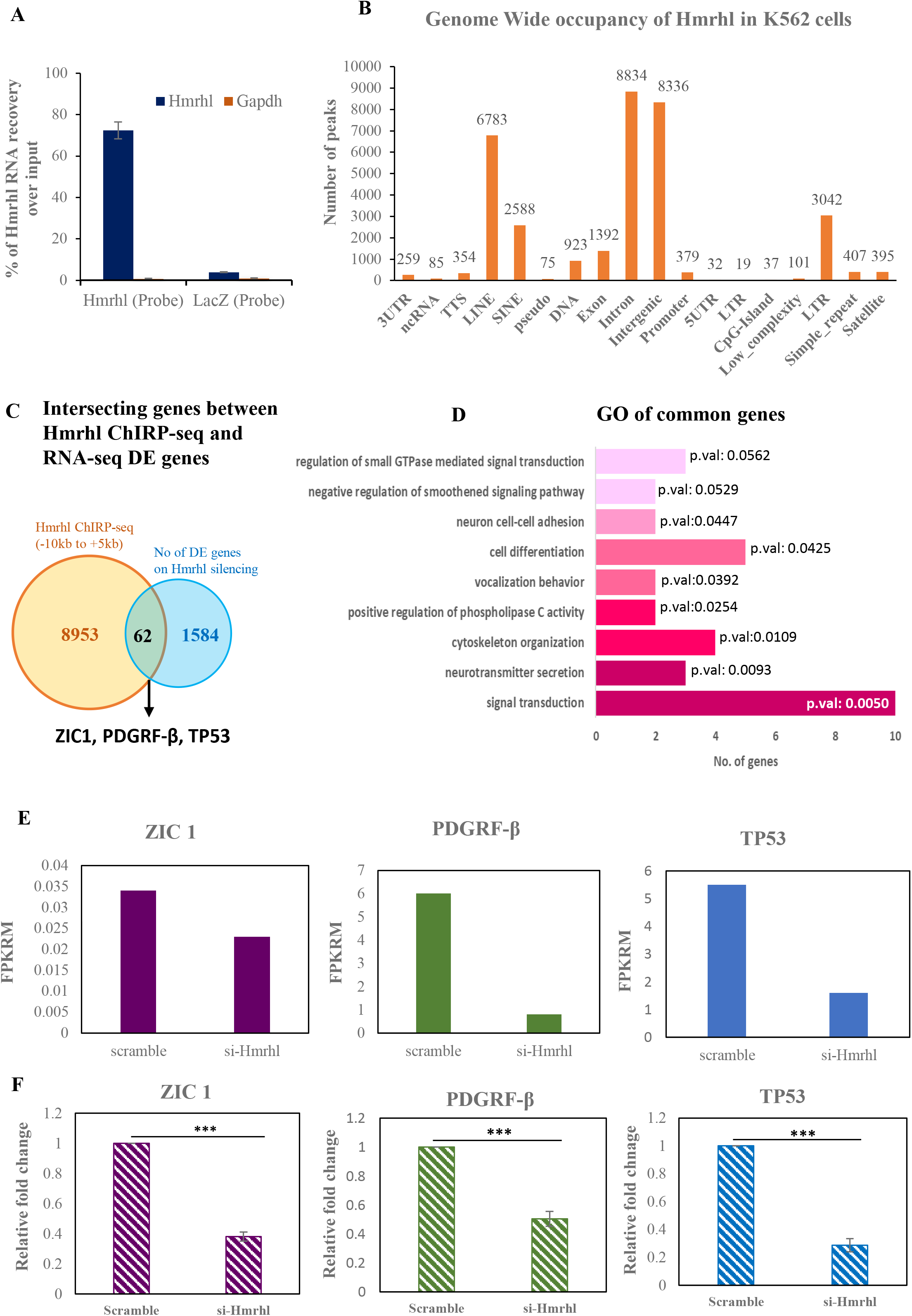
ChIRP-Seq analysis of Hmrhl in K562 cells (A) Plot showing efficiency for Hmrhl pulldown over LacZ in ChIRP experiment. (B) Annotation of peaks (C) Venn diagram representing overlapped genes obtained from intersection of RNA-seq and ChIRP-seq dataset. (D) Gene Ontology of common genes thus obtained. (E) FPKRM values depicting levels of *ZIC1*, *PDGRFβ* and *TP53* after Hmrhl knockdown as obtained from RNA-seq data analysis. (F) expression pattern of *ZIC1*, *PDGRFβ* and *TP53* determined by RT-qPCR after Hmrhl silencing in K562 cells. The data are presented as mean ± SD from three independent experiments, *p<0.05, **p<0.01, ***p<0.001 by student’s t-test

To get a better understanding of the genes which are perturbed upon Hmrhl RNA silencing because of physical association at the chromatin level, we overlapped the high throughput sequencing data from both RNA-seq and ChIRP-seq study. For this analysis we used -10 kb upstream of transcription start site (TSS) and +5 kb downstream of TSS of genes as our domain of target genes to explore the occupancy of Hmrhl RNA. With a stringent cut-off of more than 10-fold enrichment for ChIRP-seq, we obtained 62 overlapped genes between two sets of data (Figure 6C, Supplementary file 5). Gene ontology of these common genes revealed that most of them belong to biological processes like signal transduction (p-val:0.00506), cell development and differentiation (p-val: 0.0425) along with others like cellular adhesion (p-val: 0.04478) and migration (p-val: 0.0866) (Figure 6D, Table 2). These results very well corelates with our GO analysis of RNA-seq genes where signaling and developmental pathways were prominently enriched (Figure 2 D, E & F). Interestingly, the biological processes and pathways obtained after GO of common genes like signaling, cell adhesion, migration etc. are well associated with the cancer development and malignances. We further analysed the subset of common genes based on their role in signaling pathways in cancer as well as role in the developmental process and subsequently identified *PDGFRβ*, *TP53* and *ZIC1* as our top hits that were physically occupied with Hmrhl and also significantly dysregulated upon Hmrhl silencing (Figure 6C). We also calculated the FPKM value of each corresponding genes from RNA seq data, and this analysis shows that the expression of *TP53*, *PDGFRβ* and *ZIC1* are significantly altered upon Hmrhl silencing in comparison to control (Figure 6E). We further tested the expression of *PDGFRβ*, *ZIC1* and *TP53* genes under siRNA mediated silencing condition of Hmrhl by real-time PCR (Figure 6F). Our results are consistent with what we observed from RNA-seq data analysis.

**TABLE 2:** Biological pathways obtained after GO enrichment of common genes between RNA-seq and ChIRP-seq datasets. Detail description of genes under each pathway is given below.

| **Signal Transduction (GO:0007165, p.val: 0.005062523)** |  |  |  |
| --- | --- | --- | --- |
| *Gene* | *Fold change in expression (from RNAseq dataset)* | *Distance to TSS (from ChIRP-seq dataset)* | *Biological function* |
| TG | 2.56054 | -734 | hormone activity and carboxylic ester hydrolase activity. |
| FGD1 | 1.63956 | -319 | Activates CDC42. Plays a role in regulating the actin cytoskeleton and cell shape. |
| ANK2 | 1.69588 | -3310 | protein kinase binding and structural constituent of cytoskeleton |
| PDE1B | -1.77852 | -1499 | calmodulin binding and 3',5'-cyclic-AMP phosphodiesterase activity |
| NRXN2 | 1.51534 | -2035 | transmembrane signaling receptor activity and calcium channel regulator activity |
| NRXN3 | -2.08903 | -661 | cell adhesion molecule binding |
| PDGFRB | -3.39665 | -162 | regulation of many biological processes including embryonic development, angiogenesis, cell proliferation and differentiation. |
| CHN1 | -2.19144 | -194 | GTPase activator activity and ephrin receptor binding |
| PTCH2 | 4.70794 | -7374 | hedgehog receptor activity and hedgehog family protein binding. |
| HPGDS | -1.58043 | -1303 | calcium ion binding and magnesium ion binding |
| **Neurotransmitter Secretion (GO:0007269, p. val:0.009331)** |  |  |  |
| *Gene* | *Fold change in expression (from RNAseq dataset)* | *Distance to TSS (from ChIRP-seq dataset)* | *Biological function* |
| NRXN2 | 1.51534 | -2035 | transmembrane signaling receptor activity and calcium channel regulator activity |
| NRXN3 | -2.08903 | -661 | cell adhesion molecule binding |
| DGKI | -5.02254 | 8365 | NAD+ kinase activity and diacylglycerol kinase activity |
| **Cytoskeleton Organization (GO:0007010, p.val: 0.010913659)** |  |  |  |
| *Gene* | *Fold change in expression (from RNAseq dataset)* | *Distance to TSS (from ChIRP-seq dataset)* | *Biological function* |
| FGD1 | 1.63956 | -319 | Activates CDC42. Plays a role in regulating the actin cytoskeleton and cell shape. |
| ANK2 | 1.69588 | -3310 | protein kinase binding and structural constituent of cytoskeleton |
| WTIP | -2.31319 | -7255 | involved in several cellular processes such as cell fate determination, cytoskeletal organization, repression of gene transcription, cell-cell adhesion, cell differentiation, proliferation and migration |
| CTNNA2 | 1.67243 | -1990 | structural molecule activity and structural constituent of cytoskeleton |
| **Positive regulation of phospholipase C activity (GO:0010863, p. val: 0.025440324)** |  |  |  |
| *Gene* | *Fold change in expression (from RNAseq dataset)* | *Distance to TSS (from ChIRP-seq dataset)* | *Biological function* |
| PDGFRB | -3.39665 | -162 | regulation of many biological processes including embryonic development, angiogenesis, cell proliferation and differentiation. |
| PTAFR | -2.05699 | -3443 | G protein-coupled receptor activity and lipopolysaccharide binding |
| **Vocalization behavior (GO:0071625, p.val:0.039299)** |  |  |  |
| *Gene* | *Fold change in expression (from RNAseq dataset)* | *Distance to TSS (from ChIRP-seq dataset)* | *Biological function* |
| NRXN2 | 1.51534 | -2035 | transmembrane signaling receptor activity and calcium channel regulator activity |
| NRXN3 | -2.08903 | -661 | cell adhesion molecule binding |
| **Cell differentiation (GO:0030154, p.val: 0.042536642)** |  |  |  |
| *Gene* | *Fold change in expression (from RNAseq dataset)* | *Distance to TSS (from ChIRP-seq dataset)* | *Biological function* |
| IGSF10 | 1.70255 | -5900 | control of early migration of neurons expressing gonadotropin-releasing hormone |
| CPLX2 | -1.76176 | -9666 | Negatively regulates the formation of synaptic vesicle. Positively regulates a late step in exocytosis of various cytoplasmic vesicles |
| TDRD6 | -2.02059 | -4622 | involved in germ cell development |
| TP53 | -1.89693 | -148 | Acts as a tumour suppressor in many tumour types; induces growth arrest or apoptosis depending on the physiological circumstances and cell type. |
| ZIC1 | -4.16215 | 1079 | transcriptional activator. Involved in neurogenesis |
| **Neuron cell-cell adhesion (GO:0007158, p. val:0.044788079)** |  |  |  |
| *Gene* | *Fold change in expression (from RNAseq dataset)* | *Distance to TSS (from ChIRP-seq dataset)* | *Biological function* |
| NRXN2 | 1.51534 | -2035 | transmembrane signaling receptor activity and calcium channel regulator activity |
| NRXN3 | -2.08903 | -661 | cell adhesion molecule binding |
| **Negative regulation of smoothened signaling pathway (GO:0045879, p.val:0.05296437)** |  |  |  |
| *Gene* | *Fold change in expression (from RNAseq dataset)* | *Distance to TSS (from ChIRP-seq dataset)* | *Biological function* |
| SERPINE2 | -2.36671 | -3485 | signaling receptor binding and serine-type endopeptidase inhibitor activity. |
| PTCH2 | 4.70794 | -7374 | hedgehog receptor activity and hedgehog family protein binding. |
| **Regulation of small GTPase mediated signal transduction (GO:0051056, p.val:0.056180682)** |  |  |  |
| *Gene* | *Fold change in expression (from RNAseq dataset)* | *Distance to TSS (from ChIRP-seq dataset)* | *Biological function* |
| FGD1 | 1.63956 | -319 | Activates CDC42. Plays a role in regulating the actin cytoskeleton and cell shape. |
| CHN1 | -2.19144 | -194 | GTPase activator activity and ephrin receptor binding |
| ARHGA P29 | -2.32394 | -1824 | GTPase activator activity and PDZ domain binding |

These results therefore suggested that Hmrhl might have the functional role of a potential regulator of *PDGFRB*, *TP53*, and *ZIC1*. We were curious to investigate further the mechanism that facilitates lncRNA Hmrhl to select the chromatin region throughout the genome for its association. For this, we first looked for common sequence motifs enriched in the Hmrhl-bound genomic regions in our ChIRP-seq dataset and identified a strong GA-rich sequence, motif 1 (motif e-value: 1.00E-04) that accounts for 80.36% of these sequence reads and a second motif 2 (motif e-value: 1.00E-04), a CT-rich sequence which covers the rest of 24.54% (Figure 7A & B, Supplementary file 6). This indicates that GA-rich sequence plays dominating role in the targeting of Hmrhl lncRNAs across the genome. Next, we analysed the promoter region of our genes of interest (i.e. *PDGFRβ*, *TP53*, and *ZIC1*) and observed the enrichment of the ChIRP-seq peak for the presence of motifs region by FIMO in the MEMO suite (Figure 7C). Then we generated the potential triplex formation structures at those sites using the Triplexator program. Interestingly, triplex forming oligonucleotides (TFOs) thus obtained for all the three genes were also GA-enriched with high TFO score (Figure 7D, Table 3). Recently, using different techniques, several studies had validated the formation of RNA-DNA triplex structure by GA-rich homopurine sequences (100–102). These reports also established gene regulatory mechanism by lncRNA via RNA-DNA triplex formation, where the triplex acts as an anchor for the recruitment of chromatin modifiers in proximity to the gene promoters (103, 104). In our study, triplex forming oligonucleotides (TFOs) as scanned by the Triplexator software within the Hmrhl RNA detected TFOs with high score (Table 3) with overrepresentation of GA-rich sequences (Figure 7D). This raises the possibility of triplex formation by the GA-rich sequences between target genes and Hmrhl RNA and therefore we hypothesize that this might be the probable regulatory mechanism employed by Hmrhl RNA.

**TABLE 3:**
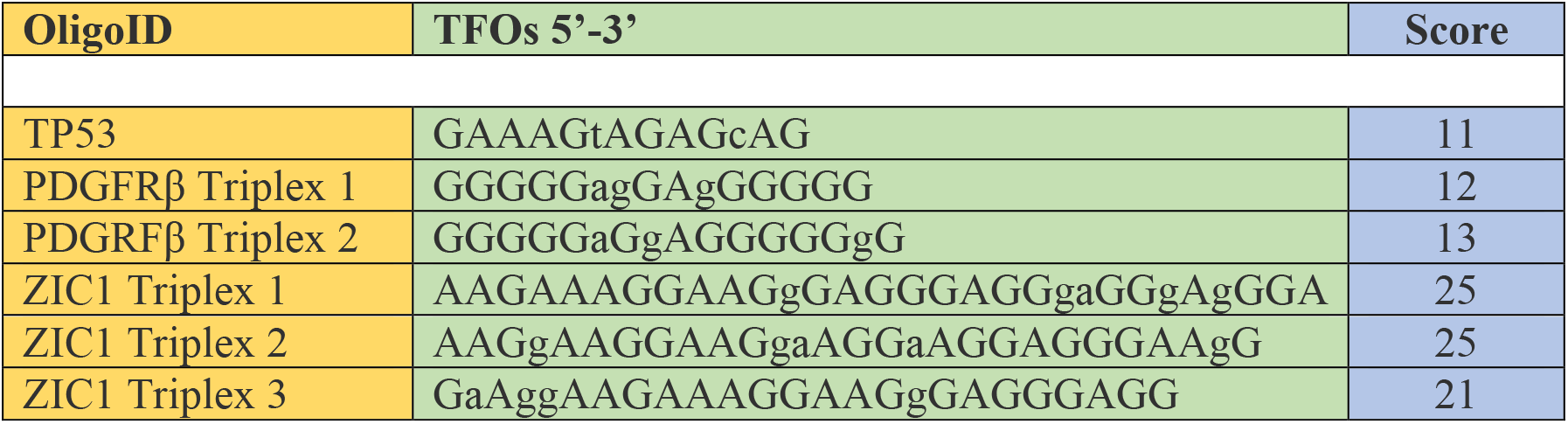
Triplex Forming Oligonucleotides (TFOs) predicted by Triplextor program

**Figure 7:**
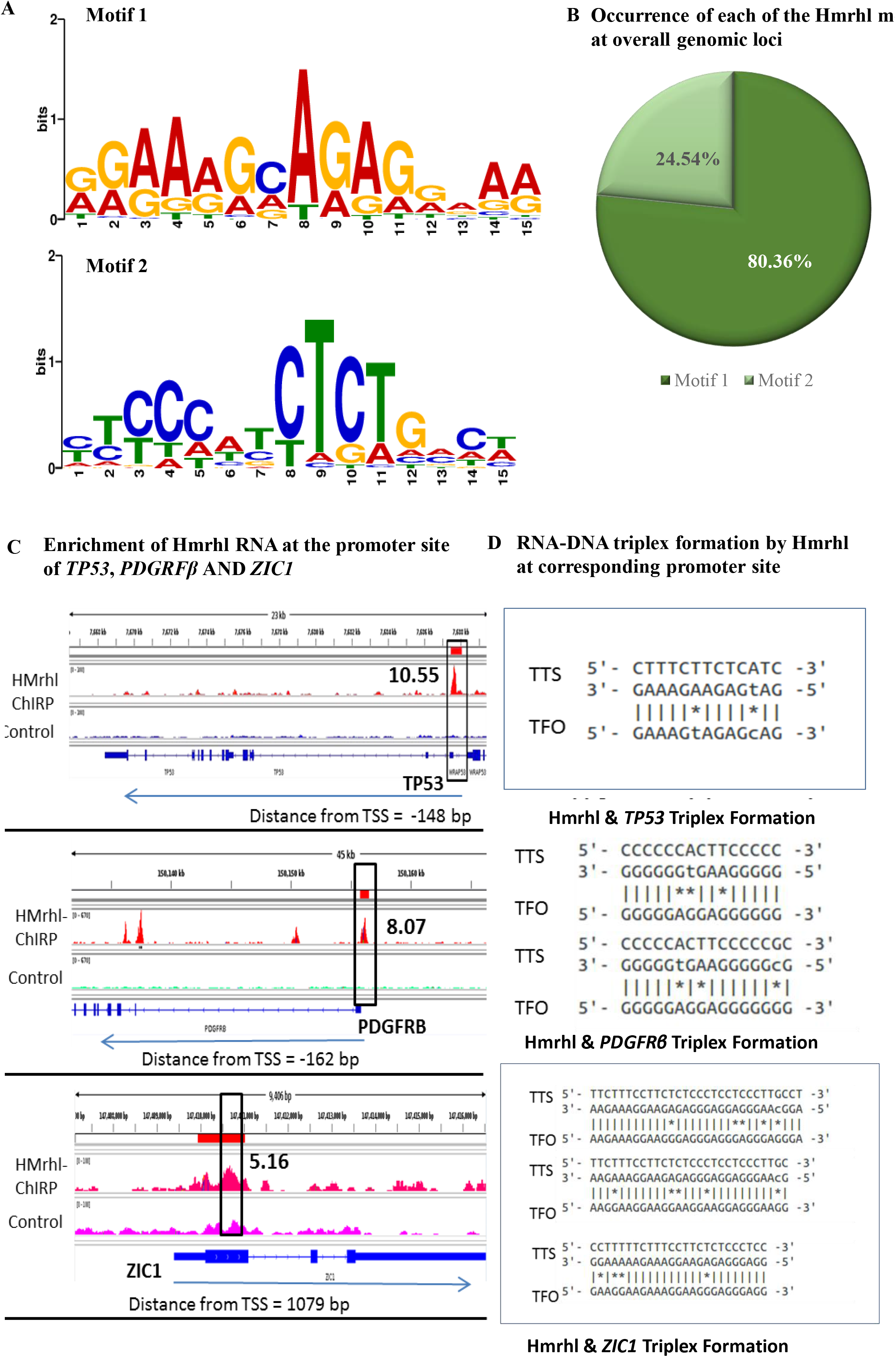
Regulation of target genes expression via Hmrhl by possible triplex formation at chromatin level. (A) Predicted motifs enriched in all Hmrhl peaks in ChIRP-seq. (B) Distribution of Motifs for genome occupancy for Hmrhl on target genes. (C) Enrichment of Hmrhl at the promoter region of *ZIC1*, *PDGRFβ* and *TP53* in ChiRP-seq analysis. (D) Predicted DN-RNA triplex formation at the target sites using Triplextor.

### TAL1 regulates Hmrhl expression in K562 cells

Since Hmrhl RNA plays a pivotal role in regulating key genes and pathways related to cancer pathobiology, we wanted to examine how Hmrhl RNA itself is regulated in K562 cells. We first tried to identify all possible TFs that has potential binding site within the promoter region (−3kb to +0.5kb) of Hmrhl with the help of GP-Miner program (Figure 8A). We found out that most of them have potential role in leukemia, hematopoiesis as well as in other type of cancers (Figure 8B). In order to determine the actual physical occupancy of various TFs, particularly in K562 cell lines, we took a look at the promoter region of Hmrhl from ENCODE project consortium, and interestingly, we came up with certain transcription factors like TAL1 and GATA2 that are highly enriched within the upstream of Hmrhl (Figure 8C). Among them, TAL 1, is a primary regulator of erythroid differentiation with an established role in genesis of hemopoietic malignancies (105). We also took advantage of the Tal1 ChIP-seq dataset in K562 available from ENCODE. Integrative Genomic Viewer (IGV) tool clearly showed that, Tal1 is indeed enriched at the promoter region of Hmrhl (Figure 8C). To further experimentally validate this, we performed Tal1 Chromatin Immunoprecipitation study (ChIP) to pull-down Tal1 associated chromatin and scored for the promoter region of Hmrhl. Our real-time PCR data clearly showed higher enrichment of Tal1 in comparison to mock control (IgG) (Figure 8D), thus confirming the data available from ChIP-seq dataset. Enrichment of Tal1 at the promoter of Hmrhl prompted us to ask the next obvious question that whether Tal1 regulates the expression of Hmrhl. To address this, we silenced *Tal1* using siRNA in K562 cell line (Figure 8E). Interestingly, we did observe an approximately 2-fold down regulation of Hmrhl after silencing *Tal1* (Figure 8F). Coherently, we parallelly evaluated *Tal1* expression upon Hmrhl down regulation and found no significant change in the expression level of *Tal1* between si-scramble and si-Hmrhl (Figure 8G & H). Thus, these observations suggest that Tal1 could be a potential regulator of Hmrhl RNA expression in K562 cells and therefore in chronic myelogenous leukemia.

**Figure 8:**
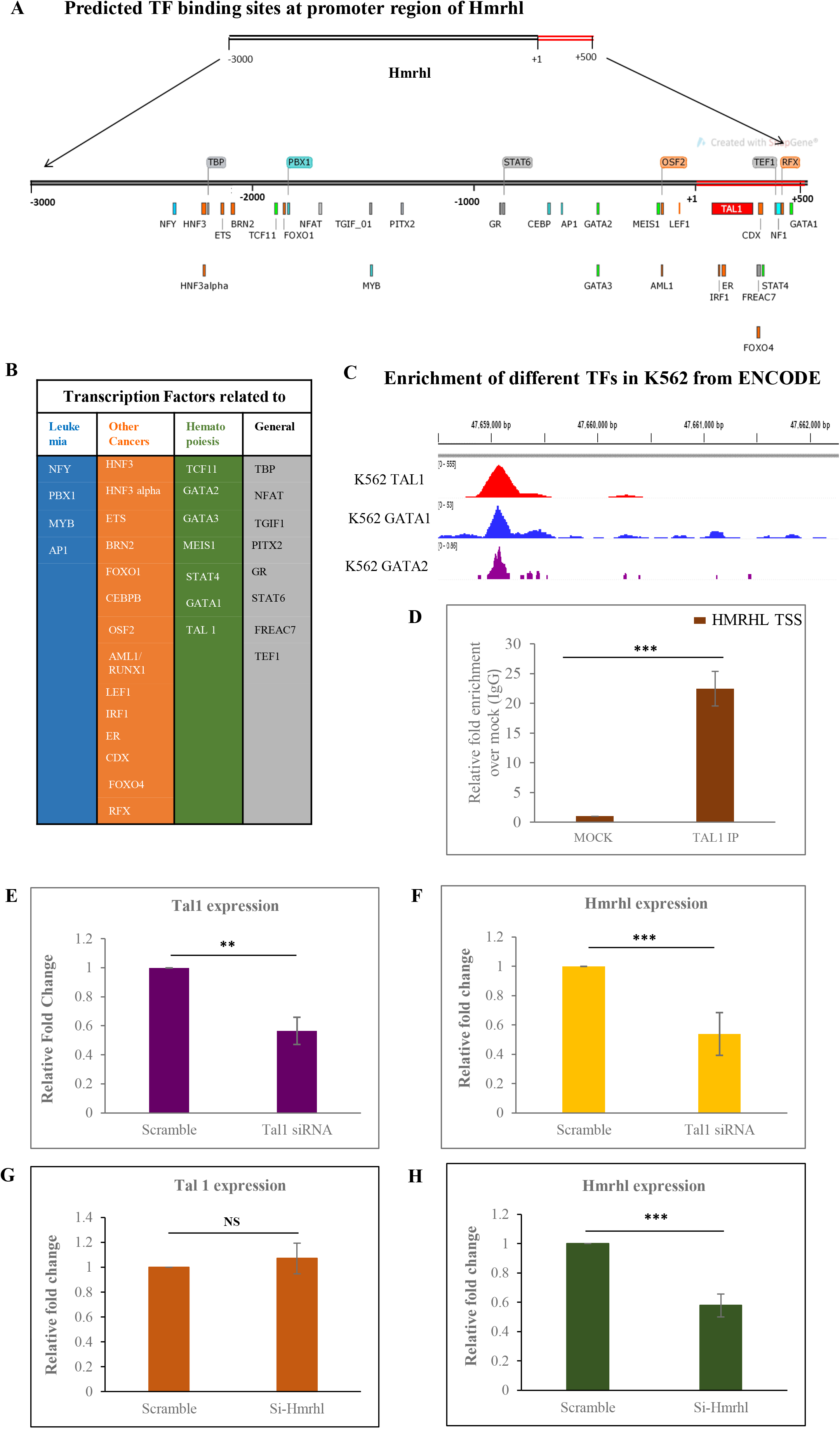
Regulation of expression of lncRNA Hmrhl by TAL1. (A) Binding location of all the possible TFs at the promoter site of Hmrhl (−3kb to +500bp) as predicted by GP Miner program. (B) Categorization of predicted TFs according to their role and involvement in leukemias, other cancers, hematopoiesis and others biological processes. (C) ChIP seq data in K562 from ENCODE establishing enrichment of few of the predicted TFs within the upstream of Hmrhl as viewed in Integrative Genomic Viewer (IGV) tool. (D) TAL1 ChIP shows high enrichment of Hmrhl promoter site when compared to mock control (IgG). (E) & (F) Expression levels of *TAL1* and Hmrhl respectively as determine by RT-qPCR after *TAL1* silencing in K562 cells. (G) & (H) evaluation of relative expression pattern of *TAL1* and Hmrhl respectively up on Hmrhl knockdown. The data are presented as mean ± SD from three independent experiments, *P < 0.05, **P < 0.01, ***P < 0.001 by student’s t-test.

## DISCUSSION

Exploring lncRNAs as novel drivers of tumorigenesis holds a strong platform and reports establishing its correlation with cancer has increased exponentially in the last decade (22, 106). Although there are experimentally supported association of 1614 human lncRNAs with various types of cancer (ln2cancer, http://www.bio-bigdata.com/lnc2cancer/statistics.html) only few of them are well documented (like H19, Xist, HOTAIR, ANRIL, NEAT1, MALAT1 etc). Most of them are not fully characterized and their functional relevance and regulatory mechanisms are still elusive (26). Following the lead from our previous study where human lncRNA Hmrhl was first reported, we tried to explore the potential of Hmrhl RNA in context of cancer phenotype in leukemia and to fully understand its regulatory targets and mechanism involved (44). Findings from the current study that Hmrhl is nuclear restricted and chromatin associated, strengthen our idea of Hmrhl RNA as a gene regulator (Figure 1C, D & E). As most of the nuclear lncRNAs associated with chromatin influence gene expression directly or indirectly by interacting with chromatin and associated proteins. For example, PINCR (p53-induced lncRNA) promotes the upregulation of a subset of p53 target genes involved in G1 arrest and apoptosis. It does so by association with the enhancer region of the candidate genes along with Matrin 3 and p53, modulating their expression upon DNA damage (107). In another example, low-irradiation-induced lncRNA, PARTICLE (promoter of MAT2A-antisense radiation-induced circulating lncRNA) represses MAT2A expression by forming a DNA–RNA triplex at the MAT2A locus, and by recruiting transcription repressive complex proteins G9a and SUZ12 (a subunit of PRC2) to the MAT2A promoter for methylation (108).

We have shown the functional relevance of Hmrhl with knockdown and transcriptome analysis using comprehensive bioinformatics approaches. This exercise revealed many interesting candidate pathways to be affected on its down regulation in K562 cell line. Inflammation mediated by chemokine and cytokines signaling and Wnt pathway been the top two hits with most number of genes to be effected (Figure 2D, Table 1). There is now overwhelming evidence to support the relationship between cancer and inflammation involving chemokines and cytokines. Predominant involvement of chemokines and cytokines in progression of cancer can be linked to their many functional capacities in metastasis, providing tumor micro-environment, survival and proliferation (109, 110). In our study also most of the dysregulated genes in this group like *CCL3*, *CCR4*, *CXCL10*, *IL-6*, *ITGB*, *ITGA*, *MYLK2*, *ReLB* are found to be associated with various cancers with their substantiated role in cell growth and proliferation, migration and metastasis (Table 1). Observations that Hmrhl promotes cell proliferation, migration and invasion in CML cell line can also be attributed to the preceded fact (Figure 5).

One interesting observation to be noticed was aberrant expression of 21 genes from Wnt signaling pathway up on Hmrhl silencing, given the fact that in our previous study no link between the two was found, as translocation of β-catenin to nucleus was not observed under the same conditions in HeK293T cells (44). It will also be fascinating to mention here that mouse counter-part of Hmrhl, mrhl was found to be inversely co-related with Wnt-signaling in mouse spermatogonial cell line, GC1-SPG while in the same species no regulation was observed between the two in mouse embryonic stem cells (42, 43). This adds to the already evident fact that functional aspect of lncRNAs are highly cellular context driven and are rarely conserved. In chronic myelogenous leukemia, Hmrhl seems to influence a completely different set of genes belonging to Wnt signaling group majority of which belongs to cadherin superfamily (eg. *DCHS1, PCDH12, CDH12, FZD9, FAT2, PCDH11Y, FRZB, SFRP5*), catenin family (eg. *CTNNA2, CTNNA3*) and signaling molecules having role in development and differentiation (Table 1). Wnt signaling pathway is crucial for the development and homeostasis of blood and immune cells. Involvement of Wnt signaling in the malignant transformation of cells has been reported earlier and its deregulation becomes apparent in the malignancies of the hematopoietic system for eg. aberrant activation of the canonical Wnt signaling is crucial for CLL pathogenesis (111). Beside affecting many important genes and TFs related to cancer, evaluation from GO analysis also shows its possible role in development and differentiation especially in neuronal development. Further exploration in this direction will be interesting as role of mrhl (mouse homolog of Hmrhl) in development of neuronal cell lineage is recently established by our group (manuscript under preparation).

Using chromatin isolation by RNA purification sequencing (ChIRP-seq) technique, it was revealed that Hmrhl RNA has efficiency to bind to a significant number of functional chromatin regions such as promoters. Integrating whole transcriptome analysis and genome-wide chromatin association of Hmrhl, we systematically identified three genes (*ZIC1*, *PDGRFβ* and *TP53*) as potential regulatory targets of Hmrhl RNA. Promoters of all of the three genes show high affinity for Hmrhl binding and a significant downregulated expression after Hmrhl depletion (Figure 6E, F & 7C, D, Table 3). *ZIC1*, which was one of the dominating TFs in our TF matrix analysis (Figure 3C, Table S2), has been reported for its oncogenic as well as tumor suppressive behavior in various cancers (112–114). *ZIC1* is also proclaimed as a one of the members of transcriptional network that controls growth arrest and differentiation in a human myeloid leukemia cell line (115, 116). Dysregulation of *ZIC1* expression may be thus associated with the outcome of our cancer phenotype study indicating Hmrhl as probable promoter of leukemogenesis with a role in cell proliferation, migration and invasion (Figure 5). As one of the most downregulated gene in si-Hmrhl RNAseq dataset, PDGRFβ was our another gene of interest to be regulated by Hmrhl. PDGFs signaling is crucial during normal development as well as has a significant role in human cancers. Stimulation of the PDGRFβ leads to the activation of the intracellular signaling pathway that further promotes cell migration, invasion, and proliferation (117, 118). Previous observation demonstrates that autocrine PDGF signaling involved in various types of malignancies, such as gliomas and leukemia (119, 120). A recent study has demonstrated metastatic and proliferative effect in murine model of pancreatic cancer induced via p53 mutation through PDGRFβ (121). This is particularly fascinating as p53 is our third gene of interest and is also mutated in K562 cell line to be translated into a non-functional protein. It will be intriguing to investigate further whether Hmrhl directly regulate *PDGRFβ* or via regulating p53 or both. As a well-known tumor suppressor, importance of p53 regulation by Hmrhl further extend its role in other cancers like breast cancer where contrary to leukemia, Hmrhl expression is highly downregulated (44).

Towards regulatory mode of Hmrhl, we postulate DNA–RNA triplex formation at the regulatory sites of target gene as a major mechanism. It is now that we know that DNA–RNA triplexes, where lncRNAs acts as a third strand is one of the mechanisms initiates/repress the expression of target gene via recruitment or stabilization of various chromatin modulators or RNA-binding proteins to the regulatory sequences (103, 122). Numerous evidences show that triplex formation at chromatin level are by motifs that are GA rich symmetrical sequences and is often considered for trans-acting lncRNA that target distant genes. For example, lncRNA MEG3 is guided to the target site via GA enriched motifs, where it facilitates the recruitment of PRC2 to the target sites (102). Other recent example includes HOTAIR lncRNA, which was found to preferentially occupy AG-rich DNA motifs across the genome. Author predicted that triplex formation by HOTAIR recruit PRC2 and LSD1 which leads to target gene silencing (104). As predicted bioinformatically, GA rich triplex formation motifs of Hmrhl RNA at target site certainly strengthen the fact of trans regulatory mechanism of lncRNA Hmrhl however, further experimental verification remains to be documented (Figure 7, Table 3). The versatility of Hmrhl function in CML needs to be explored with a thorough screening of Hmrhl-bound protein partners which only will fully reveal the its regulatory machinery through triplex formation.

We also found that expression of Hmrhl is regulated by transcription factor TAL1. TAL1 is the master regulator of haematopoiesis. It is essential for HSC renewal and commitment of haematopoietic lineages and hence its fundamental relevance in leukemogenesis evident (105). A very recent study shows that disrupting -31CBS (CTCF binding sites) relative to TSS of TAL1 cause reduce cell survival, cell cycle arrest leading to apoptosis in erythroleukemia cell line K562 (123). Enrichment of TAL1 at Hmrhl promoter and its downregulation on TAL1 depletion supports our notion of functional significance of Hmrhl in CML. Binding sites of many TFs related to haematopoiesis at the promoter region of Hmrhl certainly sparks the thought of relevant biological significance of Hmrhl in the context of haematopoiesis and erythroid differentiation.

In summation, our study provides abundant evidence to elucidate significance of lncRNA Hmrhl as an oncogene in erythroleukemia and its regulation by TAL1. Mechanistically, our data suggests its role in elevation of cell growth and migration in vitro possibly by involvement of signalling pathway genes and target gene regulation via triplex formation by binding at chromatin level. Of course, more in-depth research is required to fully comprehend the underlying mechanisms of Hmrhl RNA not only in leukemia but in other cancers which might reveal some potential therapeutic targets. Also, as indicated from the current study, involvement of Hmrhl in development and differentiation especially in neurogenesis and haematopoiesis needs further exploration to understand the complexity behind these processes.

## FUNDING

This work was supported by the Department of Biotechnology, India (BT/01/COE/07/09). M.R.S.R. acknowledges Department of Science and Technology for J. C. Bose, S.E.R.B. Distinguished fellowships and year of science chair professorship. S.R.C acknowledge S.E.R.B for National postdoctoral fellowship. S.D. acknowledges D.B.T for postdoctoral fellowship.

## ACKNOWLEDGEMENTS

We thank central facility, JNCASR. We thank Dr. Narendra Nala for assistance in flow cytometry. We acknowledge Suma B.S for her assistance with Confocal Imaging Facility.

## Notes

### Competing Interest Statement

The authors have declared no competing interest.

