## Supplemental Tables for "LncRNA Hmrhl regulates expression of cancer related genes in Chronic Myelogenous Leukemia through chromatin association"

**Table S1. Sequences of primers and probes used in this study**

| | | |
|---|---|---|
| RT-PCR Primers | HMRHL_F | TTGGCTTGGTTGGAAATGTCAGAAC |
| | HMRHL _R | ACAGGCTACAGTAGTTTGAACACAGC |
| | 18s rRNA_F | GTAACCCGTTGAACCCCATT |
| | 18s rRNA _R | CCATCCAATCGGTAGTAGCG |
| | GAPDH_F | AATCCCATCACCATCTTCCAGGAGC |
| | GAPDH_R | TGGACTCCACGACGTACTCAGC |
| | U1snRNA_F | CTTACCTGGCAGGGGAGAT |
| | U1snRNA_R | CAGTCCCCCACTACCACAA |
| | MALAT1_F | GACGGAGGTTGAGATGAAGC |
| | MALAT1_R | ATTCGGGGCTCTGTAGTCCT |
| | PDGFRB_F | CCTGCTATGAGGCTTTGGAG |
| | PDGFRB_R | GACAAATGTGCAACCACCTG |
| | ZIC1_F | TCCTACACGCATCCCAGTTC |
| | ZIC1_R | GAGGGCGATAAGGAGCTTGT |
| | p53_F | CCAGCCAAAGAAGAAACCAC |
| | p53_R | CCTCATTCAGCTCTCGGAAC |
| | Tal1_F | CTCTCGGCAGCGGGTTCTTT |
| | Tal1_R | GGGAAGGTCTCCTCTTCACTCGAT |
| TAL1 ChIP Primers | TAL1 binding site at Hmrhl TSS_F | CTTCCCGCCTCTTCACTTCACTA |
| | TAL1 binding site at Hmrhl TSS_R | TTGCTGCCATATAGAGATGACTTGC |
| ChIRP Probes | 1_HMRHL ChIRP | 5'AGTGATTTTACTATGTGTCATT 3'-Biotin-TEG modification |
| | 2_HMRHL ChIRP | 5'CTTGAGGCAAGATTTATCATCA 3'-Biotin-TEG modification |
| | 3_HMRHL ChIRP | 5'GAAAATGTCAGAGATGGGAGGG 3'-Biotin-TEG modification |
| | 4_HMRHL ChIRP | 5'ATACTGCATAGAAACAATGCCA3'-Biotin-TEG modification |
| | 5_HMRHL ChIRP | 5'GGAAAGCAGAGGAAAGTTAAGT 3'-Biotin-TEG modification |
| | 6_HMRHL ChIRP | 5'GCTTACATAGGTCTTTTATTGA 3'-Biotin-TEG modification |
| | LacZ_1 | 5'CCAGTGAATCCGTAATCATG 3'-Biotin-TEG modification |
| | LacZ_2 | 5'TCACGACGTTGTAAAACGAC 3'-Biotin-TEG modification |
| si-RNAs | Hmrhl siRNA_1_Sense | CAGUUGGAAUUGAGAUGUAUU |
| | Hmrhl siRNA_1_ Antisense | 5'-PUACAUCUCAAUUCCAACUGUU |
| | Hmrhl siRNA_2_Sense | GUGACAAAGCGUUCGGUAUUU |

| | | |
|---|---|---|
| | Hmrhl siRNA_2_ Antisense | 5'-PAUACCGAACGCUUUGUCACUU |
| | Hmrhl siRNA_3_Sense | CACUAUAAUCGCAGUCAUUUU |
| | Hmrhl siRNA_3_ Antisense | 5'-PAAUGACUGCGAUUAUAGUGUU |
| | Hmrhl siRNA_4_Sense | GAGUUAAAUGACUGGAUUCUU |
| | Hmrhl siRNA_4_ Antisense | 5'-PGAAUCCAGUCAUUUAACUCUU |
| | TAL 1 siRNA | siGENOME SMART pool, human TAL1 from Dharmacon (cat no. M-003928-00-005) |

**Table S2. ZIC1, SP8 and RELB showed maximum number of binding sites on target TFs, names of which are represented under respective column**

| | Transcription Factor Regulators (Motifs) | | |
|---|---|---|---|
| Transcription Factor Target Sites | ZIC1 | SP8 | RELB |
| | TFAP2E | TFAP2E | TP63 |
| | TP63 | ZIC1 | WTIP |
| | ZIC1 | TBX3 | ZNF341 |
| | ASCL2 | ASCL2 | HNF4G |
| | WTIP | WTIP | MAFA |
| | ZNF341 | HNF4G | RELB |
| | IRX6 | MAFA | IRF5 |
| | RELB | IRX6 | AC010522 |
| | IRF5 | RELB | |
| | CDIP1 | IRF5 | |
| | TBX10 | FEV | |
| | | CDIP1 | |
| | | TBX10 | |
| | | AC010522 | |
