## Supplemental Figures for "LncRNA Hmrhl regulates expression of cancer related genes in Chronic Myelogenous Leukemia through chromatin association"

### Figure S1

#### Gene co-expression module (Cluster 3 & 8) with no specific functional enrichment

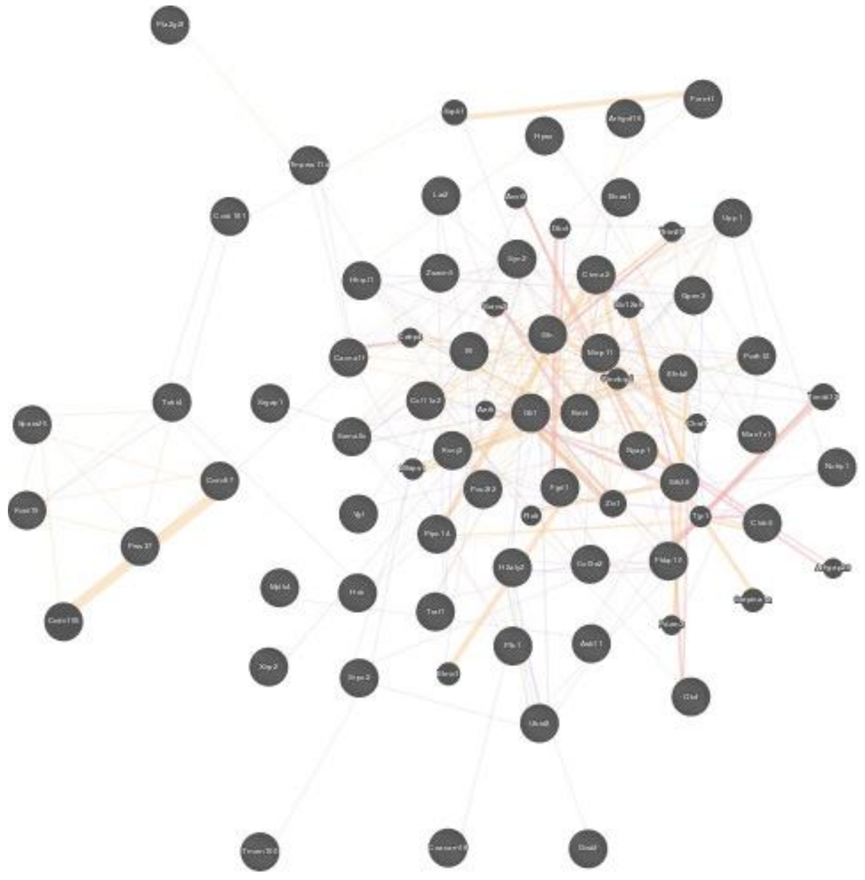

Cluster 3: Function - N/A

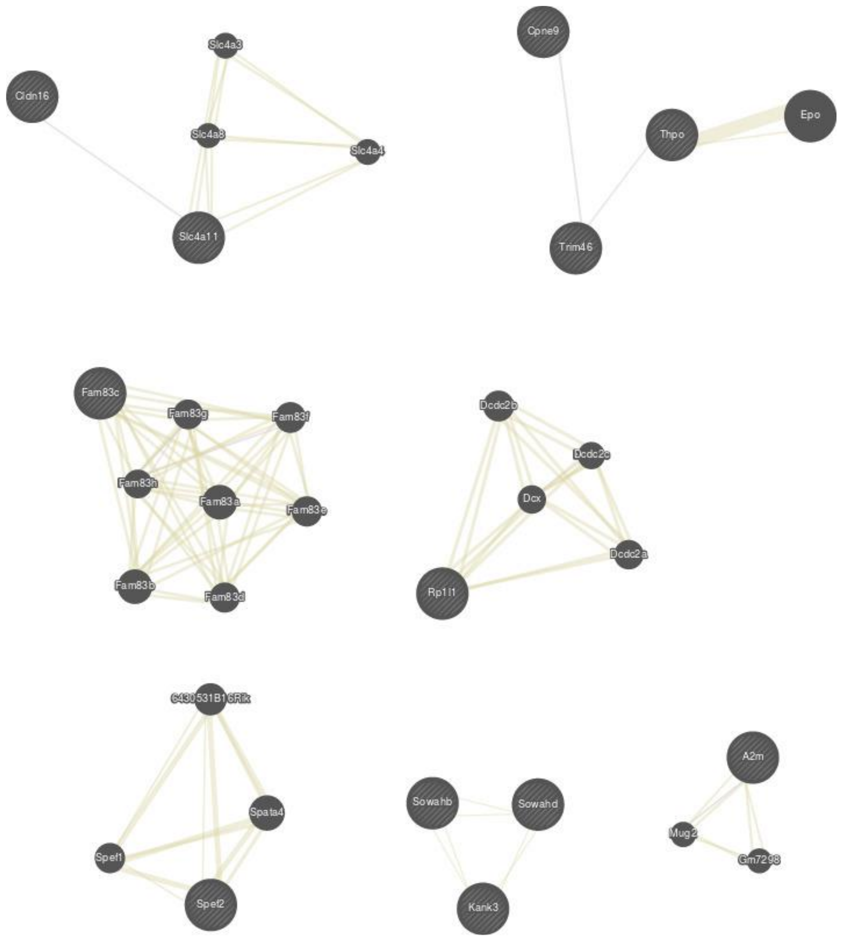


Cluster 8: Function -N/A

### Figure S2

**A** **No effect on cellular apoptosis as observed after silencing Hmrhl in K562 cells**

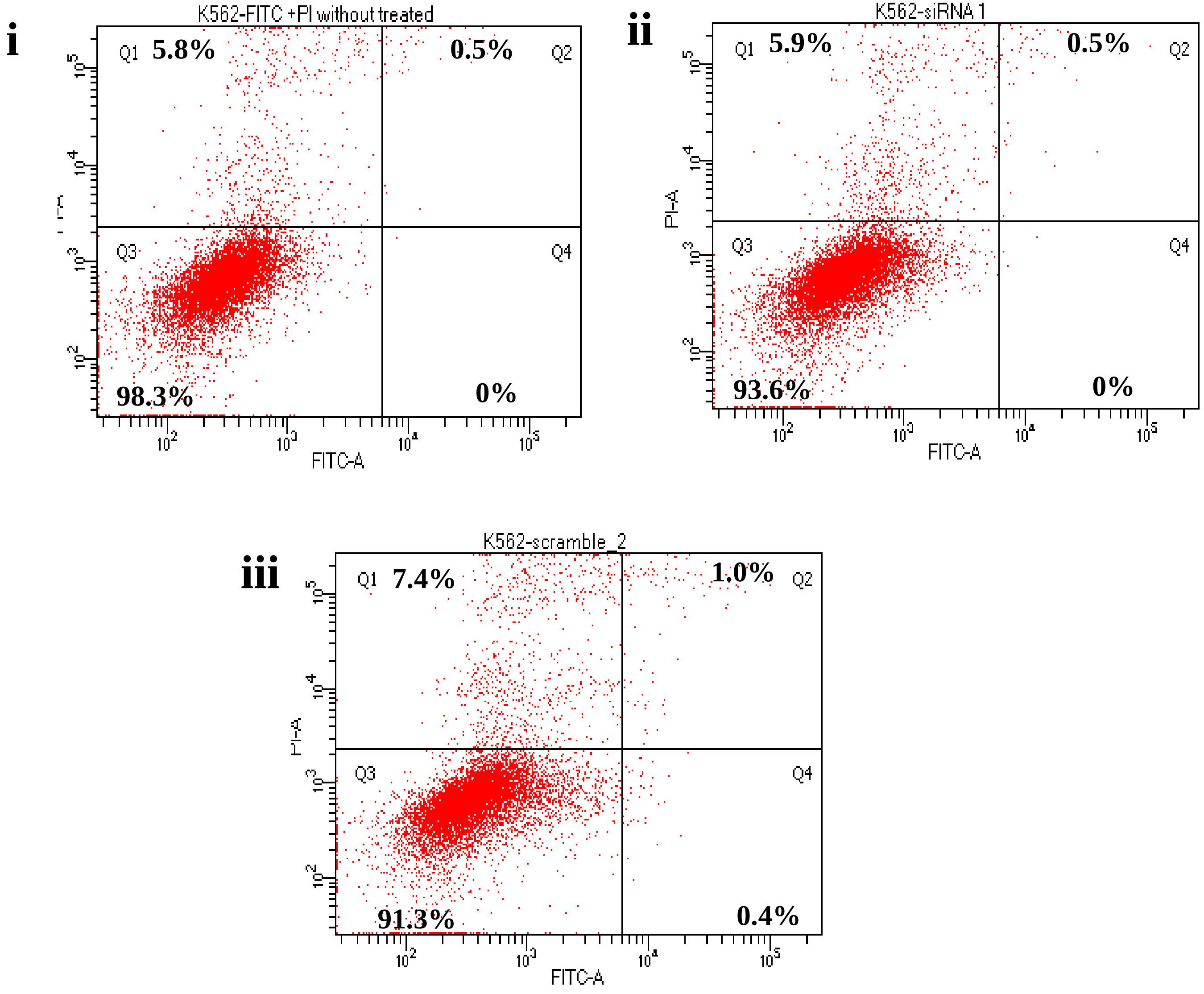


**B** **Cell Migration (CCK 8 assay)**

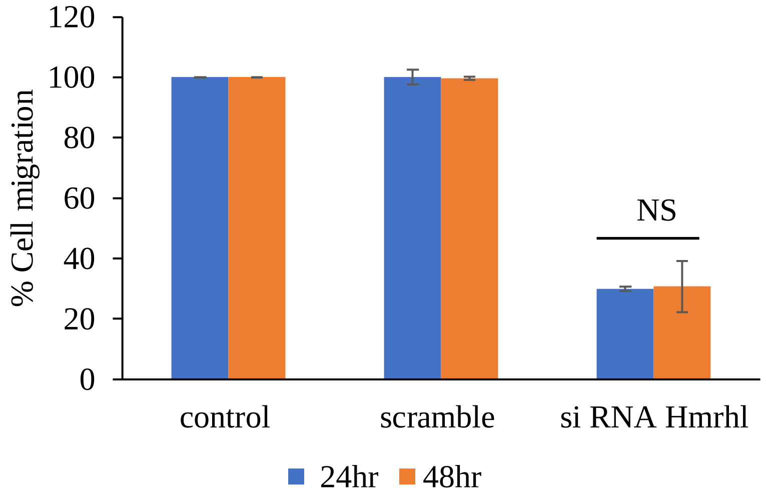


**C** **Cell Migration (Cell Count)**

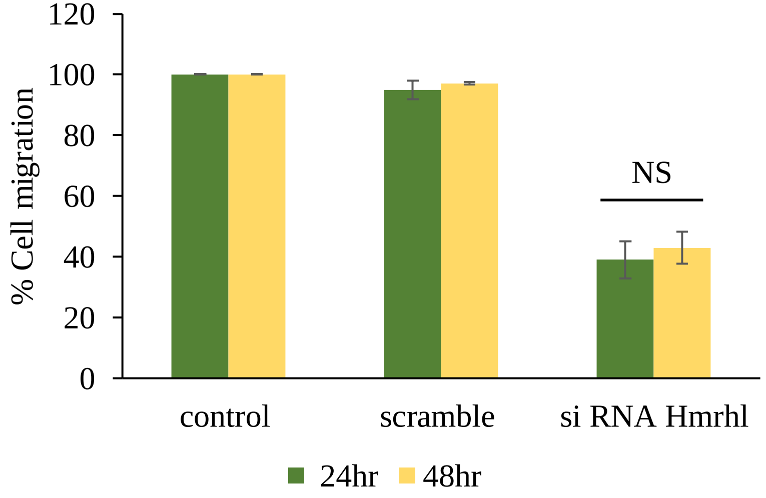
